# A conserved role of hnRNPL in regulating alternative splicing of transcriptional regulators necessary for B cell activation

**DOI:** 10.1101/2023.09.24.559201

**Authors:** Poorani Ganesh Subramani, Jennifer Fraszczak, Anne Helness, Jennifer L. Estall, Tarik Möröy, Javier M Di Noia

**Author notes:** Current address: Thermo Fisher Scientific Norway Holdings Ullernchausséen 52, 0379 Oslo, Norway (Anne Helness).

## Abstract

The multifunctional RNA-binding protein hnRNPL has been implicated in antibody class switching but its broader function in B cells is unknown. Here, we show that hnRNPL is essential for B cell activation, and thereby germinal center and antibody responses. Upon activation, hnRNPL-deficient B cells show proliferation defects and increased apoptosis. Comparative analysis of RNA-seq data from activated B cells and another 8 hnRNPL-depleted cell types reveals a common function in the MYC and E2F transcriptional programs required for proliferation, likely borne out of alternative splicing changes affecting multiple transcription regulators. Notably, while individual gene expression changes were cell type specific, several alternative splicing events affecting histone modifiers like, KDM6A, NSD2, and SIRT1, were conserved across cell types, which could contribute to gene expression changes and other phenotypes upon hnRNPL loss. In line with reduced SIRT1, hnRNPL-deficient B cells had dysfunctional mitochondria and ROS overproduction, which could contribute to defects in B cell activation. Thus, hnRNPL is essential for the resting-to-activated B cell transition by regulating transcriptional programs and metabolism, most likely through the alternative splicing of several histone modifiers.

## INTRODUCTION

Upon exposure to cognate antigen, resting B cells become activated, as a prerequisite to either differentiate into plasma cells or to enter the germinal center, where they undergo affinity-based selection to produce more effective antibodies against the infecting pathogen. The activation of resting B cells entails a switch from a quiescent, low energy spending state to a rapidly growing and proliferating state with much higher levels of transcription (Sadras *et al*, 2021; Nie *et al*, 2012; Kieffer-Kwon *et al*, 2017). To enact this change, B cells undergo rapid transcriptional and metabolic reprogramming, upregulating global transcription as well as glycolysis and oxidative phosphorylation for increased energy and biosynthetic demands (Sadras *et al*, 2021; Nie *et al*, 2012; Kieffer-Kwon *et al*, 2017). The mechanisms that coordinate transcriptional changes during the activation of B cells are thus critical for antibody responses.

RNA-binding proteins (RBPs) can modulate gene expression by affecting transcription, as well as by regulating the splicing and/or stability of the (pre)-mRNA transcripts, and thus have consequential cellular roles, including in B and other immune cells (Díaz-Muñoz & Turner, 2018; Turner & Díaz-Muñoz, 2018). Splicing factors are emerging as important regulators of B cell biology (Qureshi *et al*, 2023; Monzón-Casanova *et al*, 2018; Diaz-Muñoz *et al*, 2015; Huang *et al*, 2023). The splicing factor hnRNPL belongs to the heterogeneous nuclear ribonucleoprotein (hnRNP) family of RBPs, composed of ∼20 major members and other less abundant members, with many involved in splicing (Geuens *et al*, 2016; Han *et al*, 2010). hnRNPL is best known as an alternative splicing factor, largely regulating exon inclusion and exclusion, whereby it can prevent cryptic exon inclusion and define the usage of protein isoforms (Cole *et al*, 2015; McClory *et al*, 2018; Motta-Mena *et al*, 2010). However, several other functions have been reported for hnRNPL, some of which might be indirect. Thus, hnRNPL can regulate transcript levels, for instance by binding to and stabilizing transcripts with long 3’UTRs, protecting them from nonsense mediated decay (NMD) (Kishor *et al*, 2019). hnRNPL can also bind to several lncRNAs that can modulate gene expression (Gu *et al*, 2020), and could thus indirectly affect gene expression. A more direct role of hnRNPL in regulating transcription has been suggested in some contexts, via its interaction with the mediator subunit MED23 (Huang *et al*, 2012) or the pTEFb kinase (Giraud *et al*, 2014). Thus, it seems that hnRNPL is multifunctional and more work is required to understand its main mechanisms of action.

In line with its multifunctionality, several biological effects have been described for hnRNPL in different cellular contexts. In mouse embryonic stem cells, hnRNPL prevents apoptosis by decreasing p53 stability and localization to mitochondria (Li *et al*, 2015). Similarly, in mouse fetal liver cells, hnRNPL deletion causes apoptosis by the intrinsic and extrinsic pathways (Gaudreau *et al*, 2016). In T cells, hnRNPL deletion lead to aberrant CD45 splicing, increased proliferation and defects in cell migration (Gaudreau *et al*, 2012). hnRNPL can be overexpressed in cancer cell types, with roles in cell proliferation and migration (Gu *et al*, 2020), and promoting survival by protecting the oncogenic IgH-BCL2 fusion transcripts from NMD in B cell lymphoma (Kishor *et al*, 2019). In the CH12 mouse B cell lymphoma line, hnRNPL was shown to be required for efficient antibody isotype switching from IgM to IgA, presumably by enabling end-joining DNA repair at the end of class switch recombination (CSR) (Hu *et al*, 2015). However, hnRNPL has not been studied in normal B cells and its role during physiological CSR remains to be tested. As a cofactor of SETD2, the methyltransferase catalyzing H3K36me3, hnRNPL contributes to maintaining H3K36me3 levels (Bhattacharya *et al*, 2021b, 2021a; Yuan *et al*, 2009b). While Setd2 haploinsufficiency or knockdown in B cells confers a growth advantage and predisposes to B cell lymphoma, it does not affect CSR (Leung *et al*, 2022; Begum *et al*, 2012), suggesting Setd2-independent roles for hnRNPL at least in CSR. Other functions of hnRNPL in B cell biology and its relevance for the antibody response *in vivo* have not been studied.

The hnRNP family members exist as paralogous pairs with considerable amount of redundancy in their functions (Geuens *et al*, 2016). For example, about 50% of splicing events seem to be regulated by more than one hnRNP (Huelga *et al*, 2012). Accordingly, hnRNPL and its paralog hnRNPLL recognize CA-rich sequences and have overlapping as well as unique functions (Smith *et al*, 2013; Fei *et al*, 2017). hnRNPLL has been studied in B cells, playing an important role in plasma cell differentiation but not in B cell activation (Chang *et al*, 2015; Yabas *et al*, 2021). The RNA recognition motif of hnRNPL is also very similar to the one in PTBP1 (hnRNPI) (Geuens *et al*, 2016), which is induced and regulates alternative splicing in germinal center B cells (Monzón-Casanova *et al*, 2018). These similarities raise the question of to what extent hnRNPL, hnRNPLL, and PTBP1 might be redundant in B cells, which requires stuyding hnRNPL in B cells *in vivo*.

Here, we find that hnRNPL is essential during B cell activation. The loss of hnRNPL in B cells hinders the germinal center reaction and antibody responses. hnRNPL-deficient primary B cells undergo cell cycle arrest and apoptosis and have additional intrinsic defects in antibody class switching. In a bid to identify B cell-specific but also conserved mechanisms regulated by hnRNPL, we performed a comparative analyses of gene expression and splicing changes in mouse B cells and 8 other cell types upon hnRNPL depletion. This analysis revealed that regulating the expression of MYC and E2F transcriptional programs, as well as limiting lncRNA levels, are conserved functions of hnRNPL. Interestingly, while the effect on MYC and E2F programs is conserved across cell types, the specific gene changes are not. On the other hand, we identify conserved hnRNPL-controlled exon exclusion events in genes encoding chromatin modifiers like SIRT1, NSD2, and KDM6A. These alternative splicing events have described functional consequences on these enzymes, which could contribute to regulating gene expression and/or other hnRNPL-deficient B cell phenotypes. hnRNPL loss in B cells additionally display changes in the expression of multiple genes related to mitochondrial function. We confirm that hnRNPL is needed to maintain mitochondrial function and prevent the accumulation of ROS in resting and activated B cells, which is critical for B cell activation. Collectively, we identify both B-cell specific, as well as conserved roles for hnRNPL that provide insight into the mechanisms by which it regulates cell biology.

## RESULTS

### hnRNPL is required for the antibody response

To assess the likelihood of redundancy by paralog proteins, we first determined that *Hnrnpl* was well expressed in the mouse peripheral B cell subpopulations. In contrast, its paralog *Hnrnpll* was not expressed in the quiescent follicular (FO) and marginal zone (MZ), or in the germinal center (GC) B cells and was only expressed in plasmablast and plasma cells, albeit at 3-4-fold lower levels compared to *Hnrnpl* (**Fig. 1A**). *Ptbp1* expression was substantially higher than *Hnrnpl* in all B cell stages, but both genes were relatively similarly expressed in B cells activated *ex vivo* with T cell mimicking stimuli (**Fig. 1A**).

**Figure 1–.**
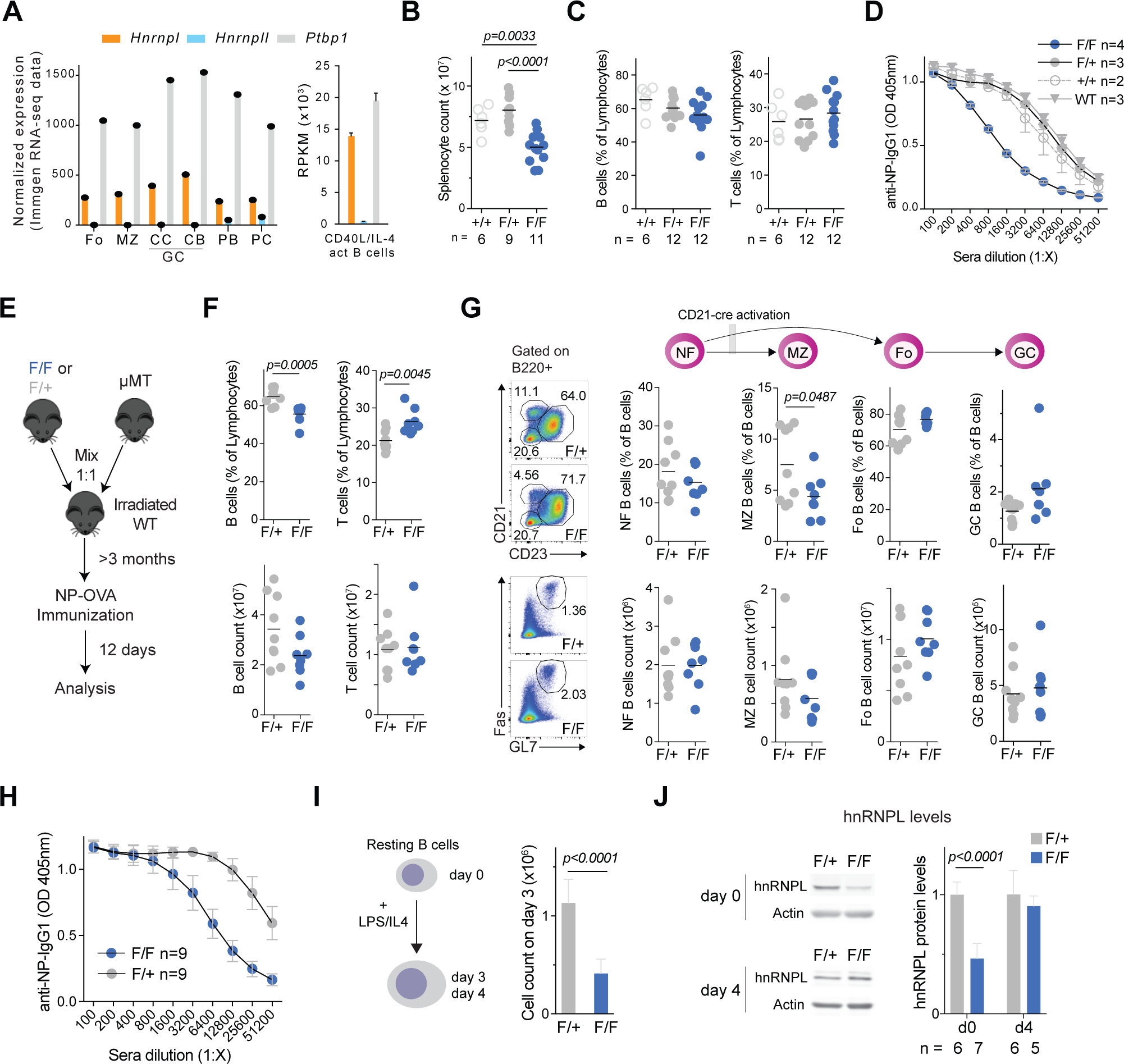
hnRNPL loss in peripheral B cells leads to a cell-intrinsic defect in antibody response. **A)** Expression levels of *Hnrnpl*, *Hnrnpll* and *Ptbp1* in mature B cell subpopulation. Follicular (Fo), marginal zone (MZ), centrocyte (CC) and centroblast (CB) germinal center (GC) B cells, plasmablasts (PB) and plasma cells (PC) from Immgen, as well as in in vitro activated B cells (mean ± SD of 4 samples, GSE120309). **B)** Splenocyte counts, **C)** splenic B and T cell proportions 11 days post-immunisation with NP-OVA, and **D)** serum anti-NP IgG1 titers 11 days post-immunisation from n *CD21-cre* (+/+), *CD21-cre Hnrnpl^F/+^* (F/+) and *CD21-cre Hnrnpl^F/F^* (F/F) and *Hnrnpl^F/+^* (WT) mice. **E)** Scheme depicting mixed bone marrow (BM) chimera experiment. BM of lethally irradiated recipient mice are reconstituted with BM cells in a 1:1 ratio from *μMT* mice and either *CD21-cre Hnrnpl^F/+^* (F/+) or *CD21-cre Hnrnpl^F/F^* (F/F) mice before immunization. **F)** Proportion and number of splenic B and T cells, **G)** newly formed (NF), MZ, FO and GC B-cell subpopulations, 12 days post-immunization with NP-OVA, and **H)** serum anti-NP IgG1 titers 11 days post-immunization from n reconstituted chimeric mice generated as in E). **I)** *Ex vivo* activation of splenic B cells purified from *CD21-cre Hnrnpl^F/+^* (F/+) and *CD21-cre Hnrnpl^F/F^*(F/F) mice and cell counts after 3 days. Results are from 9 mice each. **J)** Representative western blot and quantification of hnRNPL protein levels in B cells, either resting (d0) or 4 days post-activation as in I). All results are from at least two independent experiments. For D and H, mean ± SE OD of serial dilutions from n mice are plotted. For I and J, bars denote means + SE. P-values are indicated if differences in group means are statistically significant (p<0.05) by one-way ANOVA with post-hoc Tukey’s multiple comparison test (for B, C), unpaired two-tailed t-test with Welch’s correction (for F, G, I, J).

To assess the relevance of hnRNPL in mature B cells, we deleted hnRNPL using the *CD21-cre* system that deletes mostly in peripheral B cells (Kraus *et al*, 2004). We noticed that *CD21-cre Hnrnpl^F/F^*mice were smaller than controls (**Fig. S1A**), suggesting that *hnRNPL* was important for additional tissues in which CD21-cre excises during development (Schmidt-Supprian & Rajewsky, 2007). Accordingly, *CD21-cre Hnrnpl^F/F^* mice had smaller spleens and fewer splenocytes (**Figs. S1A, 1B**), and hence fewer total B and GC B cells (**Fig. S1B-C**). However, the proportions of total B and T (**Fig. 1C**), as well as newly formed (NF), FO, and GC B cells were not significantly affected, while MZ B cell frequency was significantly reduced (**Fig. S1D**). After immunization with the T cell-dependent antigen NP-OVA, *CD21-cre Hnrnpl^F/F^*mice produced ∼8-fold less anti-NP IgG1 than *CD21-cre* controls (**Fig. 1D**).

### An activated B cell-intrinsic role for hnRNPL supporting antibody responses

To identify B cell autonomous phenotypes of hnRNPL deficiency, we performed mixed bone marrow (BM) chimera experiments to generate mice in which only peripheral B cells have deleted *hnRNPL*. BM from *µMT* mice, which are incapable of producing B cells (Kitamura *et al*, 1991), was mixed in a 1:1 ratio with BM from either *CD21-cre Hnrnpl^F/+^* (controls) or *CD21-cre Hnrnpl^F/F^* mice and transferred intravenously into lethally irradiated WT mice (**Fig. 1E**). The body and spleen weight, as well as splenocyte numbers in chimeric mice with *CD21-cre Hnrnpl^F/F^* BM were similar to the controls (**Fig. S1E**). However, the frequency of total B cells was lower with a concurrent increase in the proportion of T cells, due primarily to a reduction in B cell numbers, while T cell numbers were not affected (**Fig. 1F**). The proportion of MZ B cells was again significantly reduced but NF, FO and GC B cells were unaffected (**Fig. 1G**). However, upon immunization with NP-OVA, chimeric mice reconstituted with *CD21-cre Hnrnpl^F/F^* BM cells showed a similar ∼8-fold defect in antibody response (**Fig. 1H**), indicating a cell-intrinsic defect caused by hnRNPL deficiency in FO B cells when responding to antigenic challenge.

To better understand the role of hnRNPL in activated B cells, we isolated resting B cells from *CD21-cre Hnrnpl^F/F^* mice and activated them with LPS + IL-4. Compared to controls, hnRNPL-deficient B cells showed a clear defect in expansion after 3 days of stimulation (**Fig. 1I**), which contrasted with the apparently normal formation of GC observed in vivo. However, while hnRNPL protein levels were reduced in the resting *CD21-cre Hnrnpl^F/F^* B cells indicating efficient *Hnrnpl* excision, activated B cells from these mice paradoxically had WT levels of hnRNPL 4 days post-stimulation (**Fig. 1J**). This indicated that cells which escaped Cre-mediated excision outgrew the Hnrnpll-null cell population over time, thus implying a major disadvantage for hnRNPL-deficient B cells in culture following activation.

We conclude that hnRNPL has a non-redundant B cell-intrinsic role in maintaining MZ B cell homeostasis and a major post-activation role in FO B cells, likely supporting cell fitness, which is required for antibody responses.

### MZ and GC B cells are hypersensitive to hnRNPL loss *in vivo*

To determine B cell fitness in vivo, we performed competitive mixed BM chimera experiments. To track hnRNPL excision in B cells, we introduced a *Rosa^mT/mG^* cre-reporter allele into *CD21-cre Hnrnpl^F/F^* mice. This allele at the *Rosa26* locus encodes constitutive expression of membrane-targeted tdTomato (mT). Upon cre recombinase expression, the mT cassette is excised, allowing for the expression of membrane GFP (mG) from a downstream cassette (Muzumdar *et al*, 2007) thus labeling cells with Cre activity and their progeny with GFP (**Fig. 2A**). BM cells from CD45.1+ WT mice were mixed with BM from either CD45.2+ *Rosa^mT/mG^ CD21-cre Hnrnpl^F/F^* or the control *Rosa^mT/mG^ CD21-cre Hnrnpl^F/+^* mice in 1:1 ratio and intravenously injected into irradiated WT recipient mice (**Fig. 2B, S2A**). Following reconstitution and immunization, the number of total B cells and the overall proportions of total B, as well as NF, FO, MZ and GC B cells were normal in both groups of mice (**Fig. S2B-C**).

**Figure 2–.**
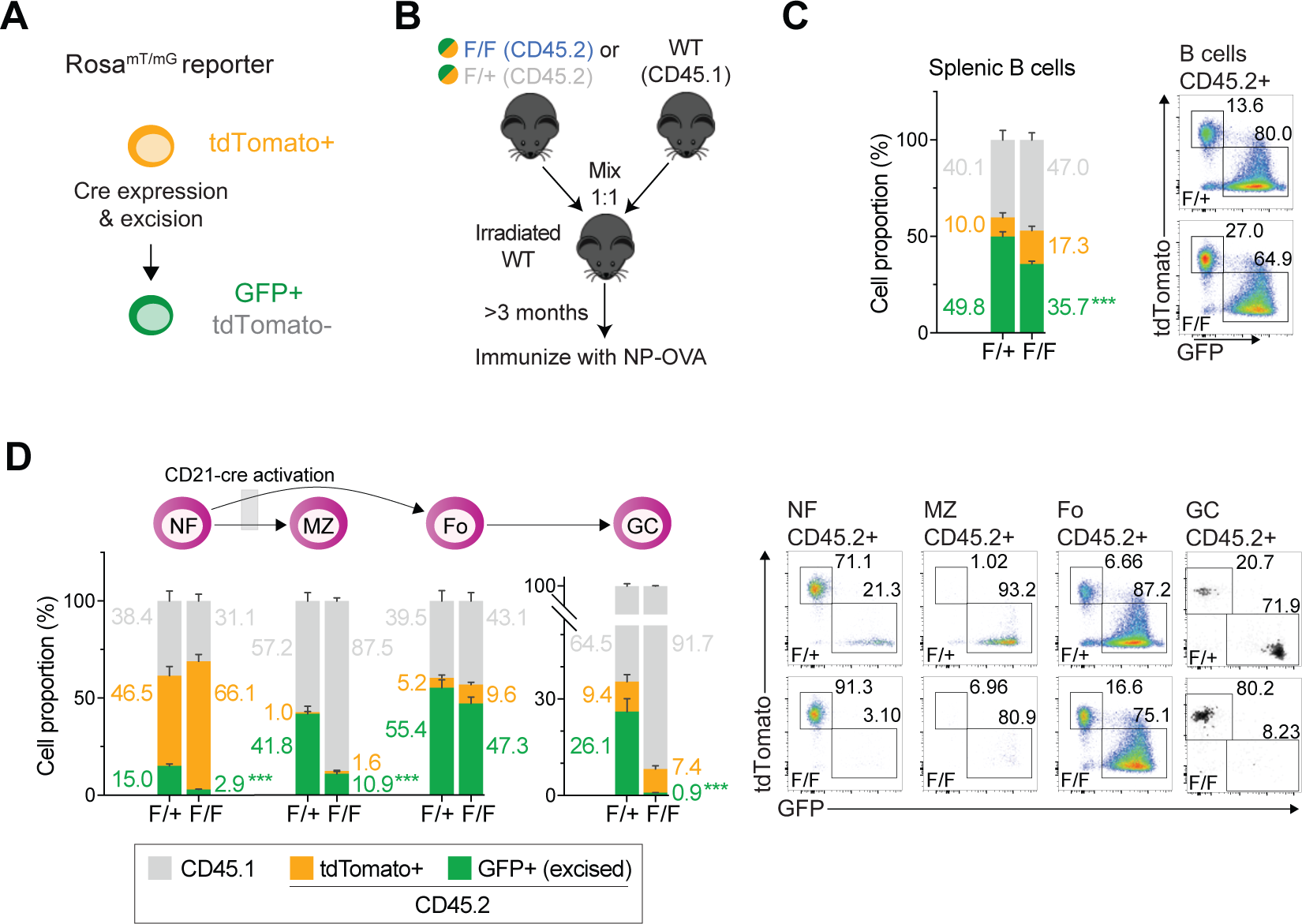
MZ and GC B cells are hypersensitive to hnRNPL loss. **A)** Scheme of Rosa^mT/mG^ reporter mouse model. **B)** Scheme of competitive BM chimera experiments. Irradiated WT recipient mice were reconstituted with CD45.1+ WT BM mixed 1:1 with BM from either CD45.2+ *Rosa^mT/mG^ CD21-cre Hnrnpl^F/+^* (F/+) or *Rosa^mT/mG^ CD21-cre Hnrnpl^F/F^* (F/F). **C)** Proportion of splenic B cells in the recipient mice originating from BM cells of CD45.1+ WT (grey) or CD45.2+ that had evidence of Cre activity (GFP+, green) or not (tdTomato+, orange), with representative flow cytometry plot of the latter analysis. **D)** as in C) for splenic NF, MZ, FO and GC B cell populations in the recipient mice. C-D) Results compiled from 2 independent experiments (8 F/+ and 8 F/F mice). Bars indicate mean ± SE. P-values are indicated if differences are statistically significant (p<0.05) by unpaired, two-tailed Mann-Whitney test. In D, *** signifies p<0.001.

While both groups of reconstituted mice showed similar proportions of CD45.1 and CD45.2 in mature B cell subsets, GFP+ cells were significantly underrepresented among B cells of the animals that received BM from *Rosa^mT/mG^ CD21-cre Hnrnpl^F/F^* mice, confirming that hnRNPL was intrinsically required for mature B cell homeostasis (**Fig. 2C**). Further examining splenic B cell subpopulations showed the onset of Cre recombinase activity after the NF B cells, as expected for *CD21-cre* (Schmidt-Supprian & Rajewsky, 2007), and no significant defect in hnRNPL-deficient FO B cells (**Fig. 2D**). In contrast, hnRNPL-deficient (GFP+) MZ and GC B cells were severely underrepresented compared to hnRNPL WT cells in the chimeric mice reconstituted with WT + *Rosa^mT/mG^ CD21-cre Hnrnpl^F/F^* BM (**Fig. 2D**). The defect was stronger in GC B cells, with even the control *Hnrnpl^F/+^* cells being outcompeted by WT cells in the control chimeras (**Fig. 2D**). Considering that MZ B cells exist in a primed or pre-activated state (Lopes-Carvalho *et al*, 2005) and GC B cells originate from activated FO cells, together with the counterselection of activated hnRNPL-deficient B cells observed *ex vivo* (**Fig. 1J**), we conclude that hnRNPL is critically required during B cell activation.

### hnRNPL is necessary for isotype switching to IgG1

Isotype switching occurs relatively early post B cell activation, and hnRNPL depletion reduces isotype switching to IgA in the CH12 mouse B cell line (Hu *et al*, 2015). To confirm this observation in primary B cells, which could be contributing to the antibody response defect, we activated B cells from the *Rosa^mT/mG^* cre-reporter mice *ex vivo* with LPS and IL-4, which allowed us to monitor isotype switching to IgG1 specifically in hnRNPL-deficient B cells. Since CSR is a stochastic process linked to proliferation (Hodgkin *et al*, 1996), we also used a tracking dye that allows enumerating cell generation span after activation. hnRNPL loss caused a severe defect in B cell proliferation (**Fig 3A** and see next section). Nonetheless, comparing cells that had undergone the same number of generations confirmed that hnRNPL loss reduced antibody class switching to IgG1 per cell division (**Fig. 3A**).

**Figure 3–.**
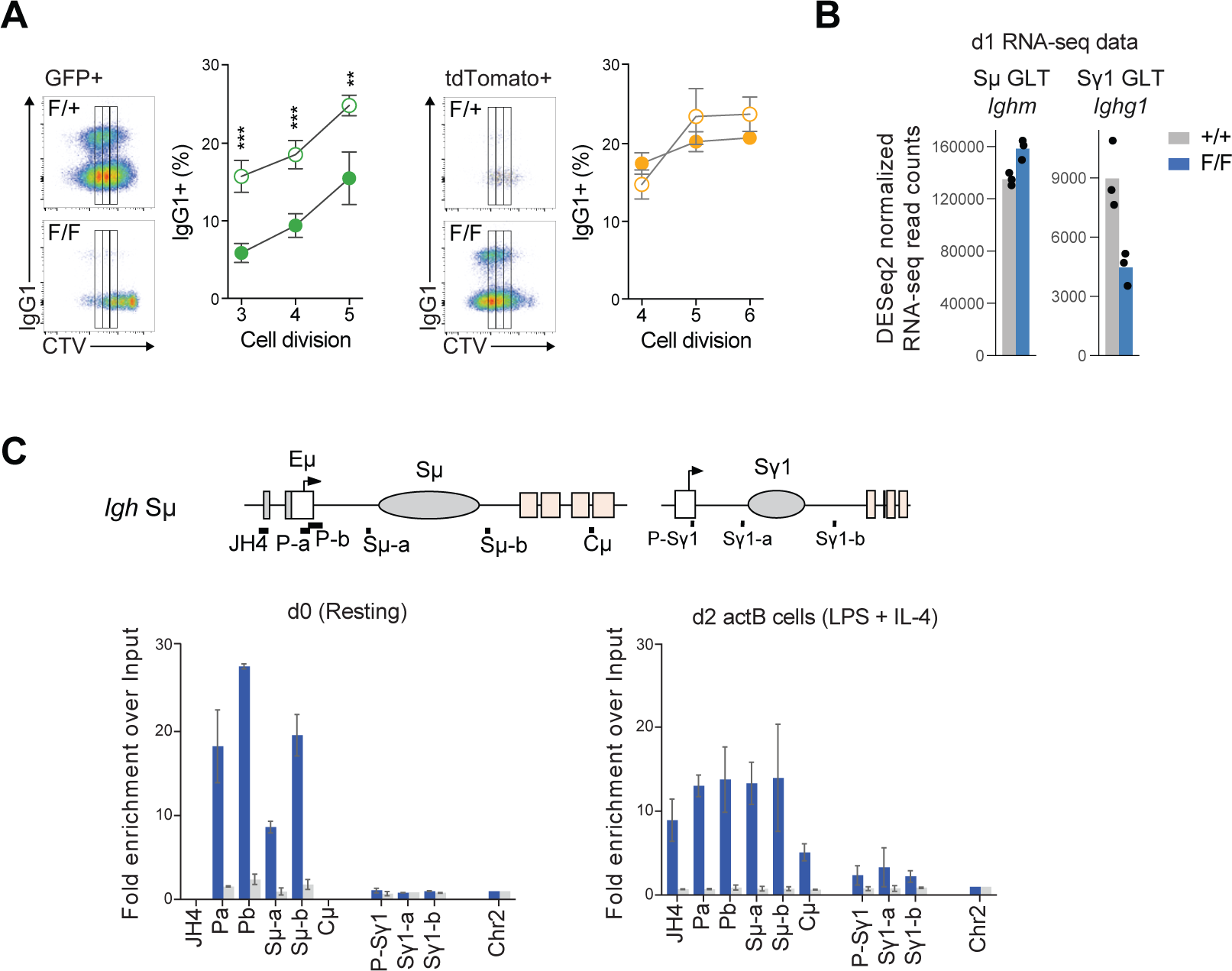
hnRNPL supports class switch recombination. **A)** Representative flow cytometry plot showing isotype switched IgG1+ B cells as a function of cell division, tracked by dilution of CTV staining in GFP+ (green; hnRNPL-excised) and tdTomato+ (orange; non-excised) B cells from *Rosa^mT/mG^ CD21-cre Hnrnpl^F/+^* (F/+) or *Rosa^mT/mG^ CD21-cre Hnrnpl^F/F^* (F/F), 4 days after activation with LPS+IL4. The plots compile mean proportion ± SD of IgG1+ cells per cell division gate as shown in the representative plots. Asterisks indicate statistically significant differences (p<0.05) by one-way ANOVA with post-hoc Tukey’s multiple comparison test (***P<0.001)**. B)** Normalised read counts from RNA-seq of Sμ and Sγ1 germline transcripts (GLTs) in WT (+/+) and hnRNPL-deficient (F/F) B cells 1 day after activation with LPS + IL-4. **C)** hnRNPL occupancy at indicated amplicons of the *IgH* locus in WT splenic B cells resting (d0) or activated with LPS + IL-4 for 2 days (d2 actB).

CSR is initiated by DNA deamination by the Activation induced deaminase (AID), which requires transcription of *IgH* Switch (S) regions, to form sterile germline transcripts (GLT) (Methot & Di Noia, 2017). RNA seq data (see next section) showed that while the donor Sµ GLT was unchanged, the acceptor Sγ1 GLT levels were reduced 2-fold in hnRNPL-deficient activated B cells (**Fig. 3B**). This contrasts with CH12 cells, in which the acceptor Sα GLT levels were not affected (Hu *et al*, 2015). However, Sα is constitutively transcribed in CH12 cells (Zhang *et al*, 2019), while acceptor GLTs in primary B cells are induced by cytokine stimuli, suggesting a potential reason for this discrepancy. To determine whether hnRNPL might be acting directly or indirectly at the *IgH* during CSR, we determined its occupancy by ChIP. hnRNPL localized primarily to the Sμ GLT region in both resting and activated WT B cells (**Fig. 3C**). At the Sγ1 region, hnRNPL was detectable only in activated B cells, when this region is transcribed, but at very low levels (**Fig 3C**). We conclude that hnRNPL associates to the *IgH*, largely at the Sµ region, and is required for CSR in primary B cells. Also, reduced Sγ1 GLT levels in hnRNPL-deficient B cells raised the possibility that hnRNPL might regulate the expression of specific genes in B cells.

### hnRNPL loss causes apoptosis and cell cycle arrest of activated B cells

To characterize the proliferation and survival defects upon activation in hnRNPL-deficient B cells, we isolated splenic B cells from the *Rosa^mT/mG^ CD21-cre Hnrnpl^F/F^* and *Rosa^mT/mG^ CD21-cre Hnrnpl^F/+^*control mice and stimulated them with LPS + IL-4. The proportion of GFP+ *Rosa^mT/mG^ CD21-cre Hnrnpl^F/F^* B cells (hereafter hnRNPL-deficient B cells) dropped starting at day 2 post-activation, and by day 4, only tdTomato+ cells (i.e., cells without hnRNPL excision) remained (**Fig. 4A, S3A**). Accordingly, hnRNPL-deficient B cells showed 2-fold more apoptosis than control cells at day 2.5 post-activation, as measured by AnnexinV staining (**Fig. 4B**), compared to heterozygous control cells. hnRNPL-deficient B cells were also smaller and less granular, indicating defective blasting following activation (**Fig. S3B**). The cell division dye staining confirmed that live hnRNPL-deficient B cells displayed a severe proliferation defect compared to control cells (**Fig. 4C**), as observed in CSR assays. This was caused by a G1 cell-cycle arrest with failure to enter S phase (**Fig. 4D**). We confirmed the effect on cell proliferation by using the induced germinal center B cells (iGB) cell culture system, in which B cells grow exponentially following activation by CD40L and BAFF displayed on the surface of feeder cells plus IL-4 (Haniuda *et al*, 2017). Cell counts, cell proliferation tracing, and cell cycle assays mirrored the results from the LPS/IL-4-activated B cells (**Fig. 4E-G**). We conclude that hnRNPL protects B cells from apoptosis and is required for cell cycle progression through the G1 checkpoint upon B cell activation.

**Figure 4–.**
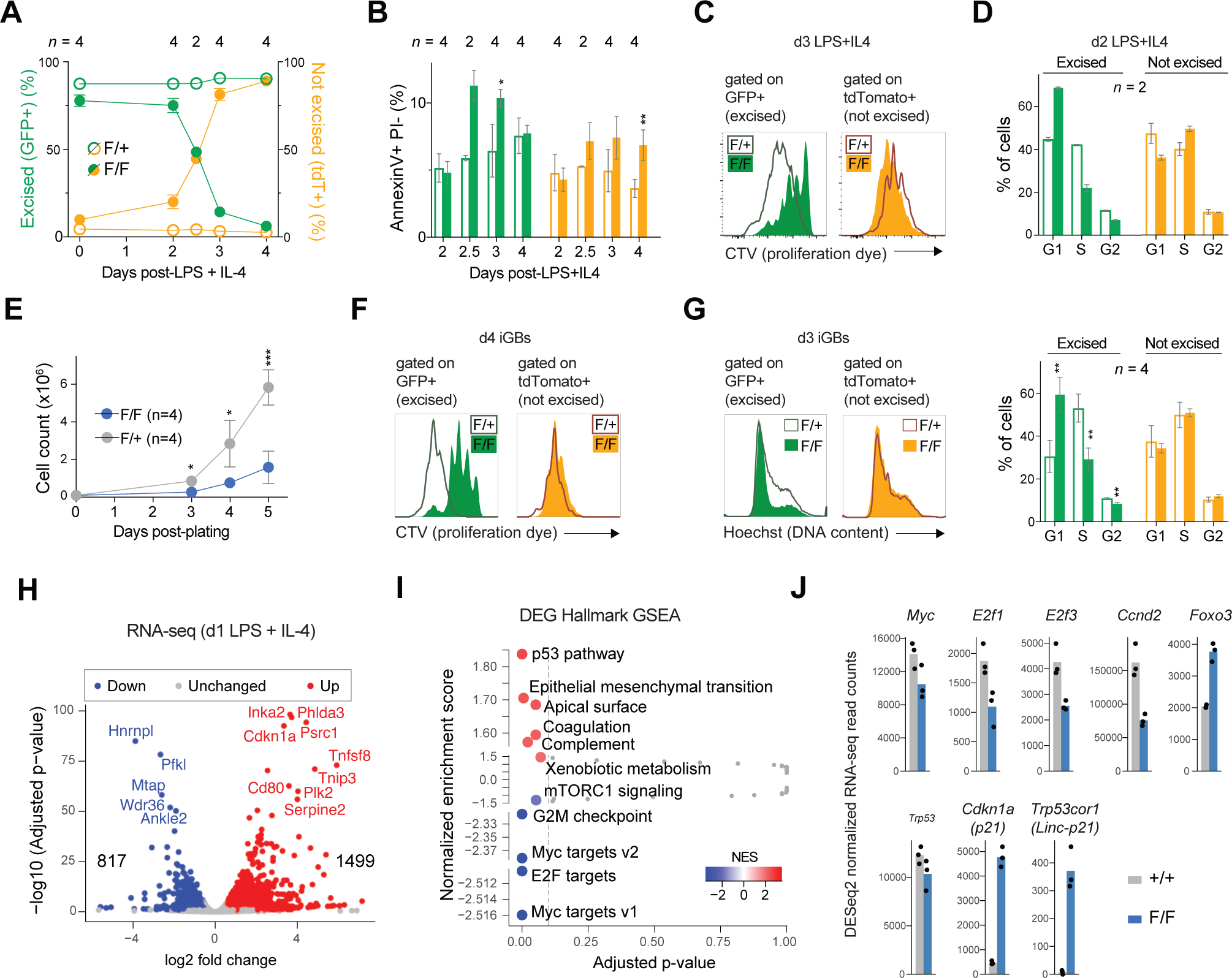
hnRNPL loss causes apoptosis and cell cycle arrest of activated B cells. **A)** Graph showing the proportions of GFP+ (green; *Hnrnpl*-excised) and tdTomato+ (orange; non-excised) B cells from *Rosa^mT/mG^ CD21-cre Hnrnpl^F/+^* (F/+) or *Rosa^mT/mG^ CD21-cre Hnrnpl^F/F^* (F/F) in culture over time after ex vivo activation with LPS and IL-4. **B)** Apoptosis, **C)** representative flow cytometry plots showing proliferation by dilution of CellTrace Violet (CTV) dye, and **D)** cell cycle profile of GFP+ and tdTomato+ B cells from F/+ and F/F mice cultured as in A). **E)** Cell count over time of induced germinal center B (iGB) cells derived from F/+ and F/F mice splenic B cells cultured on feeder cells expressing CD40L and BAFF, supplemented with IL-4 (GFP+ and tdTomato+ cells were not discriminated in this experiment). **F)** Representative flow cytometry plots showing proliferation by CTV dilution, and **G)** cell cycle profile of GFP+ and tdTomato+ iGBs from F/+ and F/F mice cultured as in E). **A-F)** Results are compiled from n mice. Symbols and bars indicate mean ± SD. Asterisks indicate statistically significant differences (p<0.05) by unpaired, two-tailed t-test with Welch’s correction (*p<0.05; ** p<0.01; *** p<0.001). **H)** RNA-seq volcano plot showing differential gene expression hnRNPL-deficient (GFP+ cells from *Rosa^mT/mG^ CD21-cre Hnrnpl^F/F^*) versus WT (GFP+ cells from *Rosa^mT/mG^ CD21-cre*) in splenic B cells activated ex vivo for 1 day with LPS + IL-4. **I)** Plot showing normalized enrichment scores for significantly enriched GSEA Hallmark terms in hnRNPL-deficient over WT cells. **J)** Normalized read counts from RNA-seq of selected genes in WT (+/+) and hnRNPL-deficient (F/F) activated B cells, symbols indicate independent RNA-seq samples, bars indicate means.

### hnRNPL sustains transcriptional programs of proliferation and dampens p53 response in B cells

To identify early and potentially causal changes that might explain defects in hnRNPL-deficient B cells, we performed RNA-sequencing on sorted GFP+ cells from *Rosa^mT/mG^ CD21-cre HnrnplF/F* and *Rosa^mT/mG^ CD21-cre hnRNPL^+/+^* controls 1 day after LPS + IL-4 activation, prior to apoptosis (**Fig. 4A,B**). Differential gene expression analysis identified 1499 up- and 817 downregulated genes in hnRNPL-deficient cells (adjusted p-value cut-off of 0.1 and ≥1.5-fold change) (**Fig. 4H**).

Functional annotation of differentially expressed genes by Gene set enrichment analyses (GSEA) using the Hallmark gene sets (Liberzon *et al*, 2015) and KEGG pathways gene ontology terms reflected cell cycle arrest, highlighting a decrease in MYC, E2F and mTORC1 signaling signatures, as well as increased p53 pathway in hnRNPL-deficient B cells (**Fig. 4I, S3C**). The combination of these changes could explain proliferation defects in these cells. Indeed, MYC and some E2F transcription factors, as well as mTORC1 signaling play key roles in cell growth, G1 to S cell cycle progression, and proliferation in activated and germinal center B cells (Nie *et al*, 2012; Patterson *et al*, 2021; Hsia *et al*, 2002; Lam *et al*, 1998). Specifically, following activation, B cells increase the expression of *Myc*, *E2f1*, *E2f3* and Cyclin D2 (*Ccnd2*) and decrease the expression of *Foxo3* to support cell cycle progression (Nie *et al*, 2012; Hsia *et al*, 2002; Lam *et al*, 1998; Hinman *et al*, 2009; Yusuf *et al*, 2004; Pae *et al*, 2021). hnRNPL-deficient B cells failed to enact these transcriptional changes (**Fig. 4J**) and showed increased FoxO signaling signatures (**Fig. S3C**). Moreover, although p53 (*Trp53*) mRNA levels did not increase, *Cdkn1a*, which is a p53 target encoding the negative cell cycle regulator and CDK inhibitor p21 (Engeland, 2018), and Linc-p21 (*Tp53cor1*), which positively regulates p21 expression (Groff *et al*, 2016), were increased more than 10-fold (**Fig. 4J**). Upregulation of *Cdkn1a* in response to activation in the hnRNPL-deficient B cells was confirmed by RT-qPCR (**Fig. S3E**). We also verified that apoptosis was a response to activation by measuring several pro- and anti-apoptotic genes in resting versus activated *CD21-cre Hnrnpl^F/F^*cells (**Fig. S3F**).

We conclude that alterations in the expression of key transcription factors and cell cycle regulators, including incomplete activation of the MYC and E2F transcriptional programs together with p53 activation, at least partly explain reduced proliferation and apoptosis of hnRNPL-deficient B cells upon activation.

### A conserved role for hnRNPL in regulating MYC and E2F transcriptional programs

Taking advantage of the availability of multiple RNA-seq datasets in hnRNPL-deficient cells, we sought to identify conserved functions of hnRNPL. Thus, we compared our RNA-seq data from activated B cells to datasets from 8 other different hnRNPL-deficient cell types: one from primary human keratinocytes (Li *et al*, 2021), five from different human cell lines (ENCODE Project Consortium, 2012; Davis *et al*, 2018; Fei *et al*, 2017; McCarthy *et al*, 2021; Bhattacharya *et al*, 2021a) and two from mouse tissues (Gaudreau *et al*, 2012, 2016). In all cell types, *Hnrnpl* expression was higher than its homolog *Hnrnpll*, with the highest ratio in mouse splenic B cells (**Fig. S4A**). In most cases, *Hnrnpll* expression did not increase to compensate for *Hnrnpl* loss (**Fig. S4A**).

To compare the biological processes affected by hnRNPL depletion in each cell type, we performed GSEA enrichment analysis for Hallmark gene sets as done for B cells. hnRNPL-deficient thymocytes exhibited much fewer differentially expressed genes than any other cell type, resulting in no Hallmark gene sets significantly enriched in these cells; but all other cell types showed several enriched gene sets (**Fig. 5A**). Remarkably, “MYC targets v1” and “E2F targets” gene sets were downregulated in 6 of the 8 informative datasets and were increased in HEK293 cells (**Fig. 5A**). For the most part, these alterations could not be attributed to downregulation of MYC or E2Fs transcripts (**Fig. S4B**). The “G2M checkpoint” gene set was also downregulated in the same cell types in which MYC and E2F targets were downregulated, consistent with a link between these programs. In contrast, most other affected gene sets were changed in only 3 to 5 datasets (**Fig. 5A**), likely reflecting context-dependent roles for hnRNPL. For example, “mTORC1 signaling” was downregulated in activated B cells and 3 other datasets, and the “p53 pathway” was increased only in mouse fetal liver and activated B cells (with the caveat that p53 activation might be impaired in some of the immortalized human cell lines). Thus, hnRNPL seems to have a conserved role in regulating the expression of MYC and E2F target genes in most cellular contexts, which at least in some cell types was independent of changes in the mTORC1 and p53 pathway, and that cannot be consistently explained by downregulation of MYC and E2F transcription factors. Another conserved feature across hnRNPL-deficient cells that emerged from our analysis was that a large proportion of the upregulated genes were long non-coding RNAs (lncRNAs), ranging from ∼45% in K562 cells to ∼20% in activated B cells, with ∼3 times more lncRNAs being upregulated than downregulated in all datasets (**Fig. 5B**) (see Discussion).

**Figure 5–.**
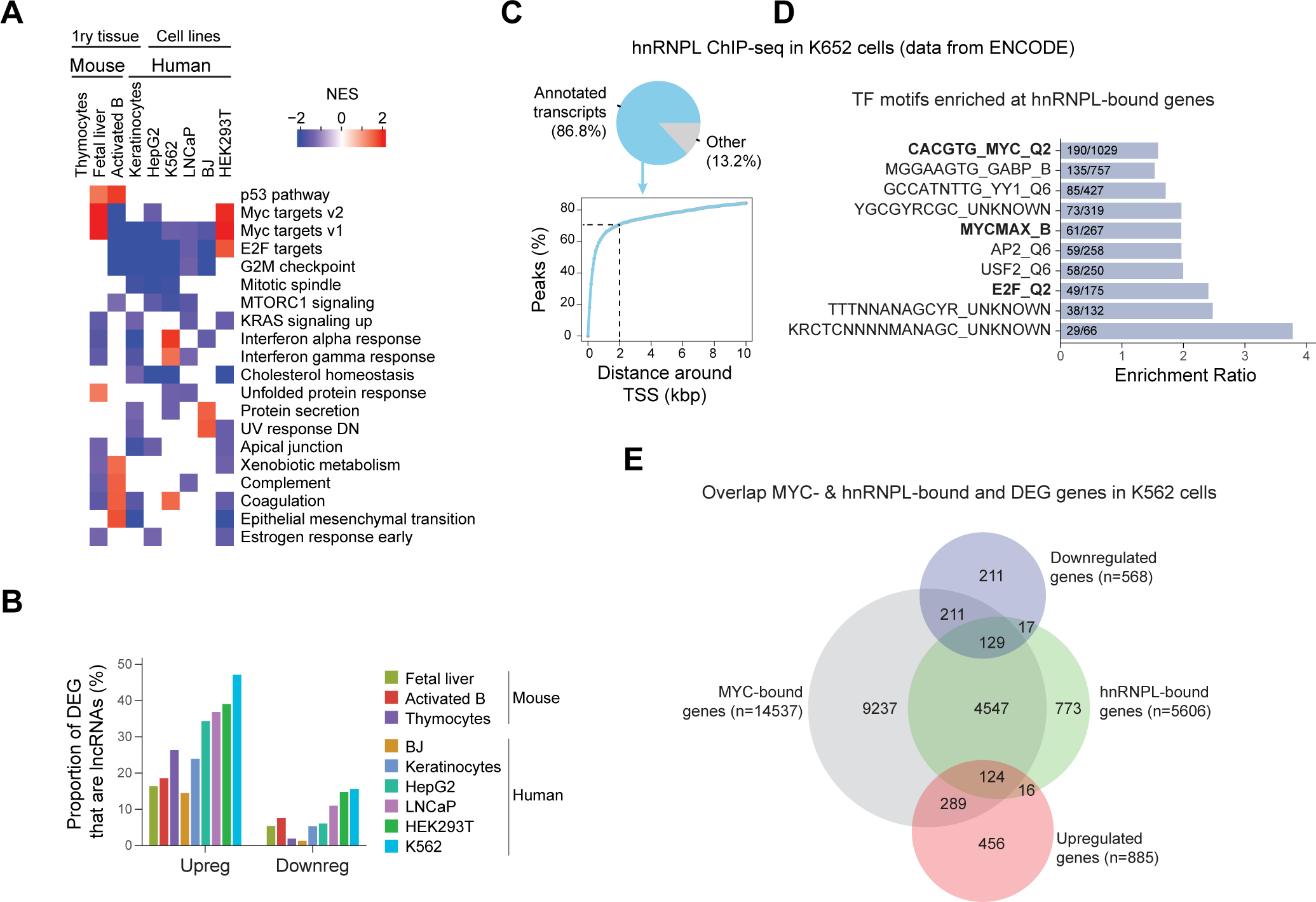
Conserved roles for hnRNPL in mouse and human cells. **A)** Comparison of GSEA analyses showing differentially enriched Hallmark gene sets for the indicated cell types. The “p53 pathway” and terms significantly enriched in at least 3 cell types were selected for display. **B)** Proportion of lncRNAs in the up- and downregulated genes in hnRNPL depleted cells. **C)** Pie chart showing the proportion of hnRNPL peaks at annotated transcripts; and cumulative distribution plot showing the distance of hnRNPL peaks from the nearest TSS, in WT K562 cells. **D)** Transcription factor (TF) motifs enriched at hnRNPL-bound gene promoters (hnRNPL peak ± 2kb from their TSS) in WT K562 cells. **E)** Venn diagrams with the overlap between MYC or hnRNPL-occupied loci in WT K562 cells, and genes significantly upregulated or downregulated in hnRNPL versus WT K562 cells.

To explore the possibility that hnRNPL could directly affect transcription, we reanalyzed hnRNPL ChIP-seq data from K562 cells, in which hnRNPL has been suggested to associate with promoters and enhancers via interactions with nascent RNAs (Xiao *et al*, 2019). Accordingly, 87% of hnRNPL ChIP-seq peaks were located within annotated transcripts, and ∼70% of those were within 2 kb of a transcription start site (TSS) (**Fig. 5C**). MYC target genes were over-represented in these hnRNPL-occupied genes, and, in fact, most hnRNPL-occupied genes were MYC targets in K562 cells (**Fig. 5D,E**). However, only a small fraction hnRNPL-occupied genes were downregulated in hnRNPL-deficient K562 cells (**Fig. 5E**). This was consistent with hnRNPL being associated to the *IgH* Sµ region in B cells, but its loss not affecting the corresponding transcript levels (**Fig 3B, C**). Similarly for a few additional B cell genes selected for being AID “off-targets” (Casellas *et al*, 2016), there was no clear correlation between hnRNPL occupancy and their transcript levels in resting and activated WT cells (**Fig. S4C**), nor were they differentially expressed in hnRNPL-deficient versus WT activated B cells (*Pax5*, *Pim1*, *Apex1* were unchanged, *Myc* was slightly reduced) (**Table S1**). Thus, without excluding that it might happen in certain instances, our analysis does not support a general role of hnRNPL in directly regulating transcription via locus occupancy.

We then looked for common differentially expressed genes (adjusted p-value < 0.2) in hnRNPL-deficient cell types. However, most of the gene expression changes upon hnRNPL depletion were cell type specific **(Fig. S4D, Table S2)**. Thus, except for a few species-specific changes, no gene was commonly up- or down-regulated in all datasets. *NUF2* and *DLGAP5* were modestly downregulated in 8 datasets (**Fig. S4D**), but they encode cytoskeleton regulators that are unlikely to regulate transcription.

We conclude that the hnRNPL has cell context-specific functions, but it is required across cell types for the maintenance of MYC- and E2F-dependent transcriptional programs, as well as to globally dampen lncRNA expression, with these effects being unrelated to any stable occupancy of the corresponding loci by hnRNPL.

### hnRNPL has a non-redundant role in regulating alternative splicing in activated B cells

Having ruled out a relationship between hnRNPL occupancy and gene expression changes in hnRNPL-deficient B cells, we turned to analyze pre-mRNA splicing.

We found 5801 differential splicing events in 2836 genes in hnRNPL-deficient versus control B cells (**Fig. 6A**). Importantly, there was minimal (367 genes) overlap between the transcripts showing significantly altered splicing events and the differentially expressed genes between hnRNPL-deficient and WT B cells (**Fig. 6B**). Concordant with hnRNPL usually regulating alternative splicing through exon inclusion/exclusion (Cole *et al*, 2015; McClory *et al*, 2018; Motta-Mena *et al*, 2010), the most common type of event affected by hnRNPL loss in B cells was exon skipping (**Fig. 6A**). Functional annotation indicated that genes whose splicing was affected by hnRNPL loss were implicated in numerous biological processes and pathways including protein degradation, autophagy, and DNA repair among the most enriched (**Table S3, Fig. 6C**). Notably, the most significantly enriched biological process was histone modification (**Fig. 6C**), which included a collection of 133 transcription factors and enzymes with well described roles in transcription regulation, such as transcription factors (*Bcl6*, *Pax5*, etc), lysine demethylases, histone deacetylases, sirtuins, and ubiquitin ligases (**Table S3** –Histone modification tab), 94 of which contained significantly changed skipped exon events.

**Figure 6–.**
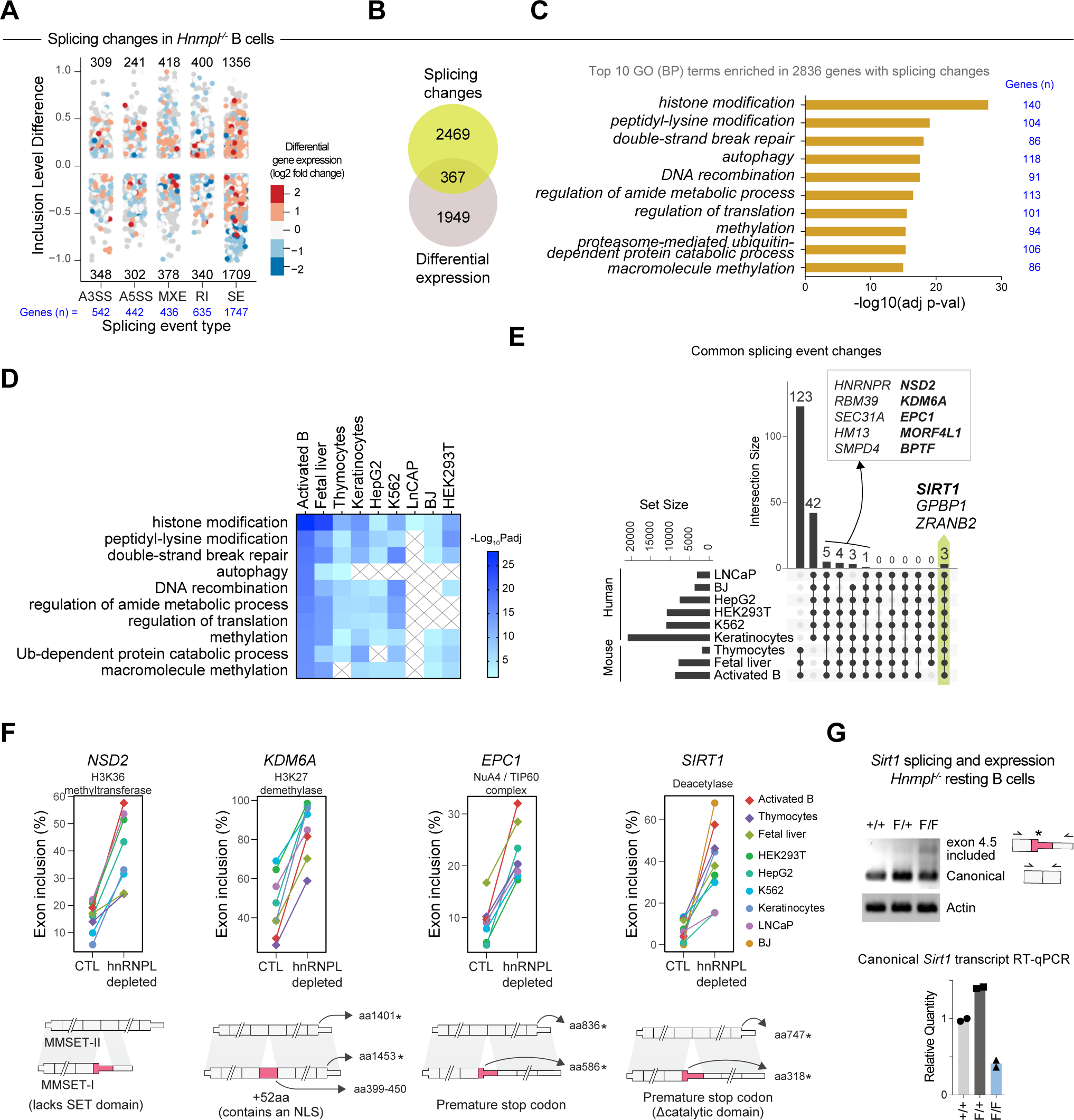
Conserved splicing events in hnRNPL-deficient cells. **A)** Splicing changes in hnRNPL-deficient (GFP+ cells from *Rosa^mT/mG^ CD21-cre Hnrnpl^F/F^*) versus WT (GFP+ cells from *Rosa^mT/mG^ CD21-cre*) activated ex vivo for 1 day with LPS + IL-4, indicating expression changes of the same genes. **B**) Venn diagram comparing the overlap between genes with significant splicing alterations and DEG in hnRNPL-deficient B cells. **C)** Top 10 enriched biological process GO terms among genes with significant splicing alterations in hnRNPL-deficient B cells. **D)** Enrichment of the indicated biological processes in genes with significantly affected splicing for each cell type. Grey X indicate the GO was not significantly enriched in the corresponding dataset. **E)** Conservation of splicing events significantly changed upon hnRNPL depletion among the indicated cell types. The inset shows the genes changed in ≥8 cell types. **F)** Exon inclusion levels of an intermediate exon in selected genes from those listed in E) in the control (CTL) and hnRNPL-depleted cell types. The schemes indicate the positions of amino acids and/or stop codons coded by the included exon in the respective human genes. **G)** RT-PCR electrophoresis gel showing the *Sirt1* canonical and the alternatively spliced transcript including the intermediate exon in resting splenic B cells from *CD21-cre* (+/+), *CD21-cre Hnrnpl^F/+^* (F/+) and *CD21-cre Hnrnpl^F/F^* (F/F) mice. Relative levels of canonical Sirt1 mRNA quantified by RT-qPCR in the same cells, are plotted below for two experiments.

We conclude that hnRNPL plays a major role in regulating alternative splicing in activated B cells that cannot be compensated by other paralogs. The splicing alterations in hnRNPL-deficient B cells seldomly affect the corresponding transcript levels directly, but a preference for regulating splicing of transcripts encoding for proteins that regulate transcription could at least partly explain the effect of hnRNPL on gene expression in B cells.

### hnRNPL has conserved roles in the splicing of transcription regulators

Given the large effect of hnRNPL deficiency on pre-mRNA splicing and its effect on transcriptional regulator transcripts in B cells, we then compared the role of hnRNPL in splicing across cell types to look for potential common mechanisms whereby hnRNPL could regulate gene expression. As in B cells, the most frequently altered type of event, and which affected the most genes, in all hnRNPL-deficient cell types was skipped exon (**Table S4**). Functional annotation of the genes with splicing affected by hnRNPL loss across cell types showed that many of the top biological processes identified in B cells were also enriched in other cell types (**Fig. 6D**). Notably, the “Histone modification” category was the only one significantly enriched in every cell type (**Fig. 6D**). We then looked for the conservation of specific splicing events across all cell types. This analysis was restricted to the exon skipping or inclusion events because there is sufficient sequence conservation to compare them (see Methods). Compared to differentially expressed genes (**Fig. S4D**), there were many more splicing changes shared between the datasets, as shown by the species-specific splicing changes (**Fig. 6E, Table S4**). Moreover, 13 differential splicing events (affecting 10 different genes) were shared in 8 of the 9 datasets, and 3 differential splicing events in *ZRANB1*, *GPBP1* and *SIRT1* genes were found in all 9 datasets (**Fig. 6E**). Interestingly, at least two of the latter can regulate gene expression; *GPBP1*, encoding a transcriptional coactivator (Hsu *et al*, 2003) and *SIRT1*, a histone and nonhistone protein deacetylase (Wang *et al*, 2019; Gan *et al*, 2020). Moreover, another 5 of the 10 genes whose splicing was changed in 8 datasets encoded for epigenetic modifiers or chromatin remodelers: *BPTF,* the NuA4/TIP60 components *MORF4L1* and *EPC1*, as well as *KDM6A* and *NSD2* (**Fig. 6E**).

Most of the conserved splicing events were increased inclusions of an intermediate exon upon hnRNPL depletion, indicating that hnRNPL normally promotes the exclusion of these exons. The consequence of each event was gene-specific, but they are all predicted to have functional consequences in hnRNPL-deficient cells, for the most part by producing different protein isoforms (**Fig 6F, S5A**). For example, hnRNPL deficiency affected the relative abundance of the *NSD2* isoforms, some encoding an H3K36 methyltransferase (Lam *et al*, 2022), produced by alternative splicing. Thus, increased inclusion of the alternative exon in hnRNPL-deficient cells biased the balance towards the MMSET-I form, which lacks the active SET domain (**Fig. 6F**) (Lam *et al*, 2022). As another example, increased exon inclusion in the *KDM6A* transcript encoding an H3K27 demethylase, introduces a nuclear localization sequence (**Fig. 6F**), which would increase its nuclear abundance (Fotouhi *et al*, 2023). In the case of *SIRT1*, the conserved hnRNPL-regulated splicing event meant the enhanced inclusion of an intermediate exon, leading to a premature stop codon that disrupts the SIRT1 deacetylase domain (**Fig. 6F, S5B**). This event was most frequent in hnRNPL-deficient activated B cells (**Fig 6F**). While not previously analyzed, we posited that this event was likely to cause nonsense-mediated decay (NMD), as suggested by reduced *SIRT1* transcript levels in 7 of the 9 cell types upon hnRNPL loss, including activated B cells (**Fig. S5C**). We then verified this result also in resting hnRNPL-deficient B cells. The transcript for the *Sirt1* form with the exon inclusion was only detected in hnRNPL-deficient B cells, as a weak amplification band consistent with it undergoing NMD (**Fig. 6G**). Accordingly, RT-qPCR showed that these cells had less canonical *Sirt1* transcript than WT cells (**Fig. 6G**). Unlike for SIRT1, the expression levels of the other genes were mostly unchanged in hnRNPL-deficient cells (**Fig. S5C**), consistent with the changes producing alternative protein forms rather than NMD.

We conclude that hnRNPL has conserved roles in splicing that affects multiple enzymes involved in transcriptional regulation, and notably specific splicing events in the pre-mRNAs of SIRT1, NSD2, KDM6A and other epigenetic and chromatin regulators, which likely underlies at least some its transcriptional effects (see Discussion).

### Mitochondrial defects in hnRNPL-negative B cells

hnRNPL loss has been associated with mitochondrial defects in fetal liver cells (Li *et al*, 2015; Gaudreau *et al*, 2016). SIRT1 is implicated in mitochondrial biogenesis and function, as well as mitophagy (Wan *et al*, 2022; Sun *et al*, 2022), partly by deacetylating PGC-1α and PGC-1β, transcriptional coactivators important for mitochondrial function (Kelly *et al*, 2009; Yi & Luo, 2010). GSEA analysis of DEG against a collection of metabolic and mitochondrial gene sets did not yield significant changes (not depicted). However, *Ppargc1b* (the gene encoding PGC-1β), which is the family member expressed in B cells, was downregulated by 50% and *Tfam*, encoding a transcription factor required for mitochondrial biogenesis (Wan *et al*, 2022), was significantly reduced in hnRNPL-deficient B cells (**Fig. S6A**). Furthermore, the leading edge of the upregulated “Xenobiotic metabolism” Hallmark gene set (**Fig. 4I**) reflected the upregulation of genes encoding enzymes that respond to redox or mitochondrial stress such as *Pink1, Cd36, Gstt2, Gstm4, Gstk1,* etc (**Fig. S6B**). Given the critical role of mitochondrial function during B cell activation (Akkaya *et al*, 2018; Boothby *et al*, 2022; Sadras *et al*, 2021; Waters *et al*, 2018), the downregulation of *Sirt1* and *Ppargc1b*, as well as indications of redox stress, prompted the analysis of mitochondrial function in hnRNPL-deficient B cells.

To measure B cell mitochondrial function, we stained resting and LPS/IL-4-stimulated B cells with two mitochondrial probes: mitotracker APC that is a measure of activity through mitochondrial membrane potential (MMP) and mitotracker green (MTG), which is incorporated into all mitochondria regardless of membrane potential and can thus be used to measure total volume/mass. hnRNPL-deficient resting B cells clearly exhibited higher MMP and MTG staining compared to controls (**Fig. 7A**). These differences were still observable 1 day after activation but equalized by day 2, possibly due to the loss of the hnRNPL-deficient cells (**Fig. 7A**). At day 2, we observed more depolarized (i.e., MTG+ MMP low) mitochondria in hnRNPL-deficient B cells (**Fig. 7A**), potentially reflecting the onset of apoptosis and/or functional defects in mitochondria. We also verified an increased gene dose ratio of mitochondrially encoded *Nd1* to nuclear encoded *Rps35a* showing increased mitochondrial DNA content in hnRNPL-deficient versus WT B cells (**Fig. 7B**), consistent with the increased mitochondrial mass (MTG signal).

**Figure 7–.**
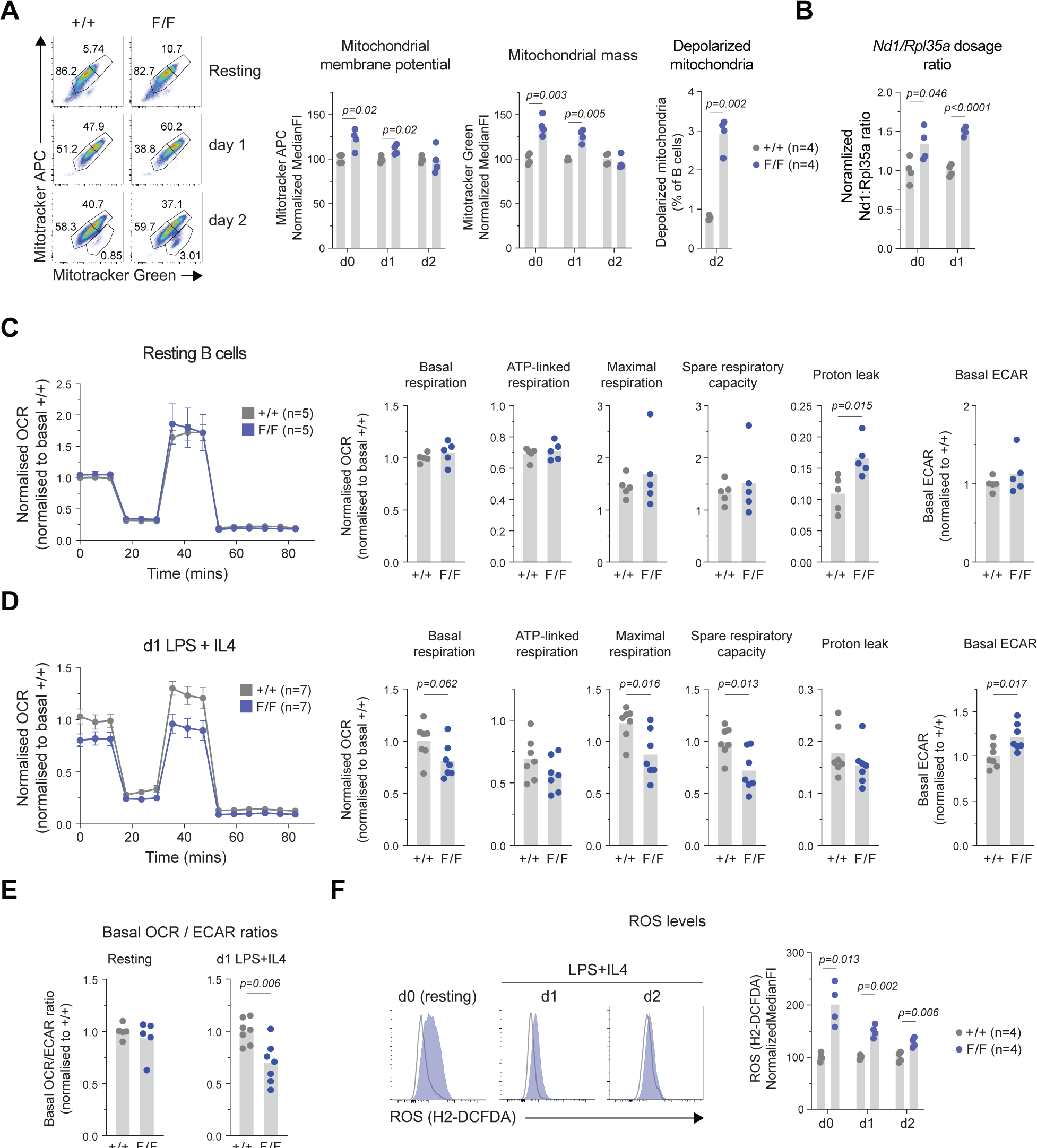
Mitochondrial defects precede B cell activation defects in hnRNPL-deficient B cells. **A)** Representative flow cytometry plots (left) and quantification summarizing normalized median fluorescence intensity of mitotracker APC and mitotracker green staining (middle) and proportion of B cells with depolarised mitochondria (right) in resting (d0) and ex vivo LPS+IL4-acitvated splenic B cells from *CD21-cre* (+/+), *CD21-cre Hnrnpl^F/+^* (F/+) and *CD21-cre Hnrnpl^F/F^* (F/F) mice followed for 2 days (d1 and d2). **B)** *Nd1:Rpl35a* gene dosage ratio calculated by RT-qPCR in cells from A). **C, D)** Mean ± SEM of normalized oxygen consumption rate (OCR) from Seahorse mitochondrial stress test, and plots showing derived parameters of mitochondrial basal, ATP-linked, maximal, spare respiratory capacity and mitochondrial proton leak, as well as ECAR measurement, in purified splenic B cells, either **C)** resting, or **D)** 1 day after activation with LPS+IL4. **E)** OCR to ECAR ratio for cells in C) and D). **F)** Representative flow cytometry plots and quantification summarizing normalized median fluorescence intensity of ROS staining in resting and B cells activated ex vivo with LPS+IL4. Plots compile results from at least 2 independent experiments. For B and C, bars and dots indicate mean ± SE. P-value summaries are indicated if differences in group means are statistically significant (p<0.05) by t-test.

Despite evidence of more mitochondria, hnRNPL-deficient resting B cells had the same basal and maximal oxygen consumption rate (OCR) compared to control cells, although they displayed increased proton leak (**Fig. 7C**). Glycolysis was unaffected in these cells, as measured by extracellular acidification rate (ECAR), indicative of lactate production (**Fig. 7C**). However, one day after activation, hnRNPL-deficient cells showed reduced coupled, maximal, and spare respiratory capacity (**Fig. 7D**), indicating mitochondrial dysfunction. In contrast, ECAR was increased, leading to a reduced OCR/ECAR ratio (**Fig. 7D,E**), suggesting that activated hnRNPL-deficient B cells were trying to compensate for an energy deficit by increasing glycolysis.

Intracellular reactive oxygen species (ROS) were substantially higher in resting and activated hnRNPL-deficient B cells, with the largest differences in resting B cells (**Fig. 7F**). Of note, ROS production preceded the observation of increased apoptosis or p53 activation (see Fig. 4B), suggesting it was not a consequence of compromised survival. The transcript levels of major ROS scavengers, electron transport chain components, and mitophagy regulators were unchanged or slightly (<1.5-fold) increased in hnRNPL-deficient B cells (**Fig. S6C**), ruling out defects in their expression. While increased ROS could simply reflect the mitochondrial mass increase in hnRNPL-deficient B cells, when coupled with defects in oxidative phosphorylation observed shortly after LPS /IL-4 treatment, increased ROS may also indicate an inherent mitochondrial dysfunction that becomes accentuated by the process of B cell activation.

We conclude that hnRNPL is required for optimal mitochondrial function in B cells, limiting ROS accumulation and supporting optimal mitochondrial respiration required during activation.

## DISCUSSION

Resting B cells are quiescent, display relatively low transcriptional activity, and are in energy deficit compared to activated B cells (Sadras *et al*, 2021; Nie *et al*, 2012; Kieffer-Kwon *et al*, 2017). Upon activation, they must rapidly increase their biomass and globally increase transcription to proliferate and function (Sadras *et al*, 2021; Nie *et al*, 2012; Kieffer-Kwon *et al*, 2017), which likely brings about requirements for increased splicing control. We show that hnRNPL is required for several aspects of the B cell activation program. Mice lacking hnRNPL in resting B cells fail to produce germinal center B cells post-activation, and have reduced MZ B cells, which exist in a primed state (Lopes-Carvalho *et al*, 2005). *Ex vivo* stimulation of hnRNPL-deficient B cells shows they fail to increase their size, progress past the G1 cell cycle phase, or proliferate, and instead they undergo apoptosis. Gene expression and metabolic changes strongly suggest a few concurrent causes for the B cell activation defect via roles of hnRNPL in regulating alternative splicing events and thereby key transcriptional programs, as well as mitochondrial function.

The number of genes affected in either their expression or splicing by hnRNPL deficiency implies that the antibody response and B cell activation defects observed are likely to have pleiotropic causes. To identify core functions of hnRNPL, we used a comparative approach that revealed that hnRNPL enables the transcriptional programs orchestrated by MYC and E2Fs in B and other cell types. MYC is a major driver of B cell activation; it is upregulated soon after activation and contributes to globally increasing transcription and metabolism to sustain cell growth and proliferation (Tesi *et al*, 2019; Caro-Maldonado *et al*, 2014; Nie *et al*, 2012; Heinzel *et al*, 2017; Kieffer-Kwon *et al*, 2017). S-phase entry further requires the activity of E2F transcription factors (Lam *et al*, 1998; Hsia *et al*, 2002; Nie *et al*, 2012; Hinman *et al*, 2009; Yusuf *et al*, 2004; Pae *et al*, 2021). Thus, suboptimal MYC and E2Fs function could largely explain the activation defect in B cells, with additional contribution of a more B cell-specific defect in mTORC signaling, as also suggested by RNA-seq. The mTORC1 complex is a metabolic sensor that enables anabolic pathways for cell growth, essential for cell cycle progression and cell growth following B cell activation (Saxton & Sabatini, 2017; Patterson *et al*, 2021; Iwata *et al*, 2016).

The mechanism by which hnRNPL regulates transcription is likely indirect. It has been proposed that hnRNPL can modulate transcription via its association with the mediator complex, or the transcriptional pause release factor pTEFb (Giraud *et al*, 2014; Huang *et al*, 2012). Indeed, hnRNPL associates with the *Trail2R* promoter and *Trail2R* is differentially regulated upon hnRNPL deletion in fetal liver cells (Gaudreau *et al*, 2016). Large scale analyses in K562 and HEPG2 cell lines also suggest and association of hnRNPL with coding genes (Xiao *et al*, 2019). We confirmed this by reanalyzing K562 data and extended by ChIP to *IgH*, *Myc* and other genes in B cells. However, our reanalysis of K562 data and of the *IgH* and other genes in B cells, fails to show a clear correlation between hnRNPL occupancy and gene expression changes upon its deletion. Thus, indirect mechanisms are more likely to explain the effect of hnRNPL on gene expression. One possibility is direct binding and protecting of a subset of transcripts from NMD, as described in other systems (Kishor *et al*, 2019). The opposite is also possible and may partly explain the conserved role of hnRNPL in limiting lncRNA expression across cell types. hnRNPL can form complexes with several lncRNAs and might regulate their abundance (Li *et al*, 2014; Klingenberg *et al*, 2018; Ruan *et al*, 2016). Changes in lncRNA abundance could also be influenced by splicing changes in multiple factors involved in non-coding RNA processing (Padj = 2.96 x 10^-12^), including the exosome subunits Exosc1, Exosc10 and Dis3 (see **Table S3**), which limits lncRNA expression (Laffleur *et al*, 2021). As lncRNAs can regulate gene expression (Li *et al*, 2014; Klingenberg *et al*, 2018; Ruan *et al*, 2016), this may further contribute to the phenotype. Our data cannot confirm or exclude these possibilities. Nonetheless, the best known and likely main function of hnRNPL is in alternative splicing by determining the inclusion or exclusion of exons in pre-mRNAs, which can alter transcript levels and protein function (Dominguez *et al*, 2018; Cole *et al*, 2015; McClory *et al*, 2018). The regulation of alternative splicing of histone modifying and other transcriptional regulators, conserved even at the level of specific splicing events for some epigenetic modifiers, emerged in our analysis as a distinct function of hnRNPL.

Controlling alternative splicing of transcription regulators provides a likely explanation for the indirect gene expression regulation by hnRNPL, which could underlie conserved and cell type-specific functions. Indeed, most conserved splicing events regulated by hnRNPL are likely to have functional consequences. For instance, the alternative KDM6 form repressed by hnRNPL would lead to higher demethylase activity in the nucleus. As described above, alternative splicing of *NSD2* pre-mRNA in the absence of hnRNPL would change the ratio of isoforms to reduce this methyltransferase activity. In addition, the overall *Nsd2* expression is ∼40% lower in hnRNPL-null than control B cells (**Fig. S5C**). Since Nsd2 contributes to the *IgH* Sγ1 expression, AID recruitment, and DNA repair in mice (Pei *et al*, 2013; Nguyen *et al*, 2017), reduced Nsd2 may contribute to the CSR defect in hnRNPL-deficient B cells. Given the association of H3K36me2 with transcription regulation (Lam *et al*, 2022), it would likely impinge on gene expression too. NSD2 deficiency reduces H3K36me2 but also H3K36me3 (Dobenecker *et al*, 2020; Hanley *et al*, 2023) and hnRNPL depletion also reduces H3K36me3 levels and transcription in cell lines (Yuan *et al*, 2009b; Huang *et al*, 2012), which could be partly regulated by its effect on NSD2 splicing. Interestingly, hnRNPL is a cofactor of SETD2, the methyltransferase catalyzing H3K36me3 onto H3K36me2. However, we can exclude a Setd2-dependent mechanism as a cause of hnRNPL-deficient B cell phenotypes because other splicing factors can compensate for hnRNPL loss in this context (Bhattacharya *et al*, 2021a, 2021b) and, in contrast to *Hnrnpl* deletion, B cell-specific deletion of *Setd2* has little effect on GC formation (Leung *et al*, 2022).

Finally, hnRNPL-deficient B cells showed reduced Sirt1 expression due to a highly conserved *Sirt1* splicing alteration likely causing NMD. Deacetylation by SIRT1 modulates the function of several histone and non-histone proteins (McBurney *et al*, 2013; Yang *et al*, 2022). SIRT1 is highly expressed in resting B cells and up to 2 days after activation (Gan *et al*, 2020) and can directly deacetylate key transcriptional regulators. Deacetylation of p53 dampens its response (Yi & Luo, 2010), of the RB protein prevents E2F inhibition to enable cell cycle progression (Wong & Weber, 2007; Jablonska *et al*, 2016; Imperatore *et al*, 2017), and of FOXO1/3a prevents it from enforcing cell cycle arrest (Motta *et al*, 2004). Deacetylation of MYC can promote or dampen its activity depending on the context (Mao *et al*, 2011; Menssen *et al*, 2012; Yuan *et al*, 2009a). Thus, reduced levels of SIRT1 in hnRNPL-null B cells could contribute to the deregulation of MYC and E2F targets, as well as p53 activation, thus hampering cell cycle progression and survival.

The importance of SIRT1 in regulating PGC-1β and overall mitochondrial function (Kelly *et al*, 2009), led us to investigate and identify mitochondrial defects in hnRNPL-null B cells. The parameters we measured indicate larger or more mitochondria, and higher membrane potential in hnRNPL-null resting B cells, yet their respiratory capacity was not increased. Activated hnRNPL-null B cells show strong indications of compromised mitochondrial respiration, and increased ECAR that would suggest an attempt to compensate via glycolysis. Elevated ROS was also evident. Since mitochondrial remodeling and respiration are critical for B cell activation, mitochondrial dysfunction most likely contributes to the activation defect of hnRNPL-null B cells. Interestingly, mice lacking canonical autophagy in B cells have similar mitochondrial defects and fewer peripheral B cells (Arnold *et al*, 2016), likely because they fail to remove damaged and aged mitochondria (Pickles *et al*, 2018). Since hnRNPL-null B cells displayed splicing alterations in genes related to autophagy and reduced Sirt1, which also regulates mitophagy (Wan *et al*, 2022; Sun *et al*, 2022), it is conceivable that they have some autophagy defect and accumulate defective mitochondria. Alternatively, B cells that fail to receive a second signal following antigenic stimulation undergo activation-induced cell death via programmed mitochondrial dysfunction, showing reduced mitochondrial respiratory capacity despite increased MTG staining (Akkaya *et al*, 2018). In any case, the elevated ROS levels linked to mitochondrial dysfunction and could cause oxidative damage and activate p53. The upregulation of p53 target genes in *Hnrnpl^-/-^* B cells is consistent with similar findings in fetal liver cells, which also show higher apoptosis, albeit this is p53-independent (Gaudreau *et al*, 2016), which we did not test in B cells. Nonetheless, the p53 activation in *Hnrnpl^-/-^* B cells likely contributes to their G1 cell cycle arrest by inducing p21 (Maillet & Pervaiz, 2012).

In conclusion, our study has revealed an essential role of the RBP hnRNPL in B cell activation and physiology and thereby in the antibody response. hnRNPL is required for cell cycle progression, preventing cell death during activation and mitochondrial function. While hnRNPL is likely to have pleiotropic effects on more than one mechanism, we provide evidence for hnRNPL functions that are essential for B cells and well conserved in other murine and human cell types, such as the regulation of MYC and E2F transcriptional programs necessary for cell cycle progression, as well a role in dampening lncRNA levels. Our data indicates that at least in part, the mechanism by which hnRNPL enables B cell activation and is essential for most cell types, is through regulating alternative splicing with a preference for pre-mRNAs encoding transcriptional regulators, whereby hnRNPL modulates the expression of many genes and critical transcriptional programs. Thus, our work suggests common underlying principles of hnRNPL function.

## METHODS

### Mice

The generation of the *Hnrnpl^F/F^* mice was previously described (Gaudreau *et al*, 2012). *CD21-cre* (Kraus *et al*, 2004) and *Rosa^mT/mG^* reporter (B6.129(Cg)-Gt(ROSA)26Sor^tm4(ACTB-tdTomato,-EGFP)Luo^/J) (Muzumdar *et al*, 2007) mice were obtained from Jackson labs (Bar Harbour, MN). Mice were housed and bred at the IRCM specific pathogens-free facility, as required to obtain the desired genotypes. All mouse work was reviewed and approved by the animal protection committee at the IRCM (protocols # 2021-1087 and 2019-05), according to the guidelines of the Canadian Council of Animal Care.

### Bone marrow chimera

Mouse BM cells were isolated by flushing the femur and tibia bones with PBS using a syringe with 23G needle. BM from *CD21-cre Hnrnpl^F/F^* or *CD21-cre Hnrnpl^F/+^* mice (CD45.2^+^), were mixed with identical numbers of BM cells from CD45.1^+^ WT mice. Lethally irradiated (9.5 Gy) C57BL6/J (CD45.2^+^) recipient mice were injected by the tail vein with 100 µl of PBS containing 5×10^6^ mixed BM cells. All recipient mice were females. Hematological reconstitution was confirmed in blood by flow cytometry 4 weeks post-injection. Mice were immunized (see below) 3 months after reconstitution. Serum was collected 11 days later, and the spleen was analyzed the next day. For mixed BM chimera experiments with tdTomato reporter, the proportion of CD45.1+ and CD45.2+ as well as GFP+ and tdTomato+ cells were determined for each B cell subset by flow cytometry. The CD45.2+/CD45.1+ ratio of all subsets were normalized to the ratio of the BM mix used to reconstitute the corresponding group of recipient mice. The proportional contribution of each genotype to each subpopulation was then calculated and plotted as a percentage.

### Immunization

2-4 months old mice were immunized intraperitoneally with 100 μg of NP_18_-OVA (Biosearch technologies) in Imject Alum adjuvant (Thermo Scientific). Mice were age- and sex-matched whenever possible depending on their availability.

### ELISA

Antigen-specific antibodies were captured from immunized mice sera in EIA High Binding surface chemistry 96-well plates (Costar) coated with NP_26_-BSA (Biosearch technologies) and incubated with dilutions of sera. Captured IgG1 was detected with Biotin-conjugated rat anti-mouse IgG1 (BD Pharmigen) followed by HRP-conjugated streptavidin (1:5000; Thermo Scientific) and developed using 2,2-azino-bis(3-ethylbenzothiazoline-6-sulfonic acid) substrate (Sigma). All antibodies are listed in **Table S5**.

### Western blotting

Cells were extracted in RIPA lysis buffer (1% NP-40, 10% glycerol, 20 mM Tris pH 8.0, 137 mM NaCl, 10% glycerol, 2 mM EDTA), containing protease and phosphatase inhibitors (Thermo Scientific) followed by sonication in a water bath for 10 mins. Extracts separated by SDS-PAGE were transferred to nitrocellulose membranes (BIO-RAD). Membranes were blocked in TBS + 5% milk and probed with primary antibodies (1h to overnight), washed 4 x 5 min in TBS + 0.1% Tween-20 and incubated with secondary antibodies conjugated to AlexaFluor680 or IRDye800 for 1 h. Signal was measured using the Odyssey CLx imaging system (LI-COR) and quantified using ImageStudiolite software. Antibodies used for WB are listed in **Table S5.**

### Flow cytometry

Lymphocytes were obtained by mashing the mouse spleen through a 70 µm cell strainer with a syringe plunger. Cells suspensions were washed in PBS and resuspended in 1mL of red blood cell lysis (155 mM NH_4_Cl, 10mM KHCO_3_, 0.1mM EDTA) for 5 min at room temperature. After washing, cells were resuspended in PBS + 1% BSA and stained with the different combinations of antibodies indicated in the results. Antibodies are listed in **Table S5**. Live cells were distinguished using DAPI or propidium iodide (PI) staining, as appropriate.

### Primary B cell cultures

Naïve primary B cells were purified from splenocytes by CD43^+^ cell depletion using anti-CD43 microbeads (Miltenyi, cat# 130-049-801) and an autoMACS (Miltenyi), or by using the EasySep™ Mouse B Cell Isolation Kit (Stem Cell, Cat. #19854) and the column-free magnet EasyEights™ (Stem Cell, Cat. #18103), following manufacturer instructions. Resting B cells were cultured at 37°C with 5% (vol vol^-1^) CO_2_ in RPMI 1640 media (Wisent), supplemented with 10% fetal bovine serum (Wisent), 1% penicillin/streptomycin (Wisent), 0.1 mM 2-mercaptoethanol (Bioshop), 10 mM HEPES, 1 mM sodium pyruvate and plated at 0.5 million cells/ml and stimulated with lipopolysaccharide (LPS) (5 μg/mL, Sigma) + IL-4 (5 ng/mL, PeproTech). Alternatively, naïve B cells were seeded onto 40LB feeder cells (a kind gift from Dr Daisuke Kitamura) (Nojima *et al*, 2011) to generate iGB cells. One day before B cell plating, 40LB cells were irradiated (120 Gy) and plated at 0.3 x 10^6^ cells per well in 2 mL (6-well plate) or 0.13×10^6^ cells per well (24-well plate) in 0.5 mL DMEM media supplemented with 10% fetal bovine serum (Wisent) and 1% penicillin/streptomycin (Wisent). Purified naïve B cells were plated on 40LB feeders at 10^5^ cells per well in 4 mL of iGB media (6-well plate), or 2 x 10^4^ cells per well in 1 mL (24-well plate), supplemented with 1 ng/mL IL-4. At day 3 post-plating, the same volume of fresh media was added to the wells, supplemented with 1 ng/mL IL-4 (PeproTech). On subsequent days, half of the volume per well was removed and replaced with media as above.

### Monitoring apoptosis, cell cycle, proliferation and isotype switching

To assess apoptosis *ex vivo*, 3 x 10^5^ cells were stained with 3 μL Annexin V-APC (cat #550474, BD Pharmigen) in 100 μL of the provided Binding buffer (1x) for 15 min at RT. Then 400 μL of Binding buffer (1x) and 5 μL of propidium iodide (20 μg/mL) were added prior to flow cytometry acquisition. For cell cycle analysis, live cells were stained with 10 μg/ml Hoechst 33342 (Thermo) in iGB media for 1h at 37C. The cell cycle stages were then calculated using FlowJo software. To assess cell division, naïve B cells were stained with 2.5μM CellTrace^TM^ Violet (Invitrogen, cat. no. C34557) in PBS for 20min at 37C before quenching with media as per the manufacturer’s protocol, just prior to their plating with cytokines or onto 40LB cells. CTV-loaded B cells were stimulated with LPS + IL-4 with conditions mentioned in the previous section to induce IgG1 switching, and analyzed after 4 days by flow cytometry.

### Reverse transcription and quantitative PCR

RNA was isolated using TRIzol (Life Technologies) or TRI-reagent (Molecular Research Center, Inc), following manufacturer’s instruction, and quantified by NanoDrop (Thermo Fisher). cDNA was synthesized from 1 μg of RNA using the ProtoScript^TM^ M-MuLV Taq RT-PCR kit and random primers (New England BioLabs). Quantitative PCR using SYBR select master mix (Applied Biosystems) was performed and analyzed in a ViiA^TM^ 7 machine and software (Life technologies). All oligonucleotides are listed in **Table S6**.

### Chromatin immunoprecipitation

The detection of hnRNPL by ChIP was performed as described previously (Rashkovan *et al*, 2014; Helness *et al*, 2021) with the addition of Disuccinimidyl glutarate (DSG) crosslinker (Thermo). Briefly, 20 x 10^6^ resting or activated B cells cells/ChIP freshly prepared from spleens were treated with 1.5mM DSG for 17min at RT. Cells were then cross-linked with 0.25% formaldehyde for resting B cells or 1% formaldehyde for activated B cells for 8 minutes and quenched with 125 mM glycine. Cells were lysed and chromatin was sonicated (Covaris E220 sonicator) to a size range of 200–500 bp. Samples were immunoprecipitated with 5 μg anti-hnRNP L antibody (Aviva ARP40368_P050. Lot: QC9464-42964) or control anti-IgG antibody (sc-2025, Santa Cruz Biotechnology) coupled with Dynabeads beads coated with Protein G (Life Technologies). ChIP DNA was analyzed by qPCR. The relative enrichment over input was calculated using the ΔΔCt method and normalized to a negative control intergenic region per sample.

### Mitochondrial analysis

For Mitotracker stainings, 500,000 cells were stained with Mitotracker Green and Mitotracker Deep Red (Thermo) at a final concentration of 20 nM each for 30 mins at 37°C in 100 μl RPMI without phenol red (Thermo) + 1% FBS. For measuring ROS levels, 250,000 cells were stained with 1 μM CM-H2DCFDA (Thermo) in 250 μl RPMI without phenol red (Thermo) + 1% FBS for 25 min at 37°C.

### Oxygen rate consumption analysis

0.5 million B cells were cultured in assay media (XF base medium (Agilent), 200 mg glucose, 0,5 mM sodium pyruvate and 2 mM glutamine, pH 7.4) for 1 h without CO_2_ prior to measurement of O_2_ consumption and extracellular acidification rate (ECAR) by XF^e^24 (Seahorse Bioscience) with sequential addition of 1 μM of oligomycin, 1 μM of FCCP and 0,5 μM of rotenone.

### RNA-sequencing

GFP+ cells from *Rosa^mT/mG^ CD21-cre Hnrnpl^F/F^*(hnRNPL-deficient) and *Rosa^mT/mG^ CD21-cre* (control) mice were FACS sorted from splenic B cell cultures, 1 day after activation with LPS+IL4. RNA was extracted using Qiagen RNeasy Mini kit. mRNA was enriched using NEBNext Poly(A) mRNA Magnetic Isolation Module (NEB) and library was prepared using KAPA RNA HyperPrep kit (Roche Diagnostics), as per manufacturer’s instructions. Equimolar libraries were sequenced on Illumina NovaSeq 6000 at the McGill University and Génome Québec Innovation Centre to generate 100 bp paired-end reads.

### RNA-seq analysis

Sequencing adapters were trimmed with fastp (0.23.1) (Chen *et al*, 2018) before alignment with STAR (2.7.9a) (Dobin *et al*, 2013) to mm10 reference genome. Gene expression counts were determined with featureCounts (Rsubread 2.12.2) (Liao *et al*, 2014) in a strand specific fashion. Differentially expressed genes were determined with DESeq2 (version 1.38.2) (Anders & Huber, 2010). Mapped reads were filtered for mapping quality (MAPQ>=1) with using samtools (1.16.1) (Danecek *et al*, 2021) to generate bigwig files using bamCoverage and bamCompare from deepTools 3.5.0 with a bin size of 1bp. Data is deposited at GEO under accession number GSE242069.

Publicly available RNA-seq datasets that were re-analyzed were as follows: primary human keratinocytes (GSE162546) (Li *et al*, 2021), HEK293T (GSE151296) (Bhattacharya *et al*, 2021a), HepG2 (GSE87985 and GSE88069) (ENCODE Project Consortium, 2012), K562 (GSE87973 and GSE88364) (ENCODE Project Consortium, 2012), LNCaP (GSE72844) (Fei *et al*, 2017), BJ fibroblasts (GSE154148) (McCarthy *et al*, 2021), mouse thymocytes (GSE33306) (Gaudreau *et al*, 2012) and mouse fetal liver cells (GSE57875) (Gaudreau *et al*, 2016). Raw FASTQ files of these datasets were downloaded with fasterq-dump tool from sra-toolkit (version 3.0.0) (https://github.com/ncbi/sra-tools) and RNA-seq and splicing analyses were performed using the same bioinformatics pipeline as done for B cells.

### Splicing analysis

For optimal detection of splice variants, multi-sample 2-pass mapping strategy was employed during read mapping, following the instructions from STAR manual (Dobin *et al*, 2013). Differential splicing analysis was performed with rMATS (version 4.2.2) with options “--variable-read-length --novelSS”. To compare human and mouse splice variants, an algorithm was developed. First, a master table of all differential splicing events across all mouse and human samples with FDR<0.2 cutoff was made. Then for each homologous gene, each unique skipped exon (SE) splicing event (defined by the start and end positions of 3 exons: the spliced exon and its adjacent exons 3’ and 5’ of it, as identified in the rMATS output tables) in mouse was compared to all splicing events in human for its homologous gene. Briefly, the sequence of each type of exon (“current”, 3’ or 5’) was obtained using the BSgenome R package (version 1.66.3; https://bioconductor.org/packages/BSgenome; Human sequences from BSgenome.Hsapiens.UCSC.hg38 version 1.4.5 and mouse sequences from BSgenome.Mmusculus.UCSC.mm10 version 1.4.3). These sequences were compared to all unique exons of the same type, across species in their homologous counterpart, using the pairwiseAlignment function from Biostrings R package (version 2.66.0; https://bioconductor.org/packages/Biostrings). A percentage identity of at least 70% for each exon type was required to warrant further consideration. The mouse and human splicing events with the highest alignment score were considered equivalent splicing events. This process was repeated after removing this set of equivalent splicing events, to look for other equivalent splicing events within the same gene. GenomicRanges package (version 1.50.2) was used to handle genomic coordinates information (Lawrence *et al*, 2013). Finally, the details of splicing events detailed in the comparative splicing analysis figure were manually checked using UCSC genome browser as well as ExPasy translate tool (https://web.expasy.org/translate) to verify splice variants and stop codons.

The sashimi plot for SIRT1 splicing event was generated using rmats2sashimiplot (https://github.com/Xinglab/rmats2sashimiplot) with the option “--intron_s 15” to scale down intron lengths. UpSetR package (1.4.0) was used to make upset plots (Conway *et al*, 2017). To normalize gene expression values across samples, qsmooth function from qsmooth package (1.14.0) (Hicks *et al*, 2018) was used. Other plots were made using ggplot2 R package (Wickham, 2016).

### ChIP-seq analysis

K562 ChIP-seq dataset is from GSE120104 (ENCODE Project Consortium, 2012). For hnRNPL and Myc ChIP-seq peaks, IDR thresholded peaks files (ENCFF854WAP and ENCFF608CXN, respectively) were obtained from ENCODE portal. For comparison with up- and downregulated genes, genes were considered Myc- and hnRNPL-bound, if a peak overlapped ± 2kb of the gene transcription start site (TSS). Transcription Factor motif enrichment analysis was performed using WebGestalt (Wang *et al*, 2017). Genome ontology and distance to nearest TSS were computed using HOMER (Heinz *et al*, 2010).

### Functional annotation analyses

KEGG pathway enrichment analysis was performed with shinyGO (version 0.77) (Ge *et al*, 2020). Gene ontology (GO) term enrichment analysis was performed with the g:GOSt module of gprofiler (Reimand *et al*, 2016). For both KEGG pathway and GO term enrichment analysis, the set size was limited from 20 to 500 elements and statistical cut-off was set to ≤0.05 (FDR or adjusted p-value, respectively). Hallmark GSEA analysis was performed using msigdbr (7.5.1) (Liberzon *et al*, 2015) and fgsea (1.24.0) (Korotkevich *et al*, 2021) R packages, with the FDR cut-off ≤0.1. The GSEA comparison heatmap was plotted with ComplexHeatmap (2.14.0) (Gu *et al*, 2016) R package.

### Statistics

The statistical tests were performed using the Graphpad Prism software version 7 or 8. The tests and significance cut-off used are indicated in the corresponding figure legends and text.

## ACKNOWLEDGEMENTS

We thank Anne-Marie Patenaude for assistance with some of the mouse work. We thank the technical assistance of E. Massicotte and J. Lord with flow cytometry; O. Neyret, S. Boissel, M. Rondeau, P. Gingras-Gélinas and F. Couderc with Sanger and NGS experiments; V. Calderon with bioinformatics; M. Laprise, E-L. Thivierge, S. Demontigny, M-C. Lavallee, C. Dube and P. Bergeron with animal technical help. Computations were made on the supercomputer Narval from École de technologie supérieure, managed by Calcul Québec and Compute Canada. The operation of this supercomputer is funded by the Canada Foundation for Innovation (CFI), Ministère de l’Économie, des Sciences et de l’Innovation du Québec (MESI) and le Fonds de recherche du Québec – Nature et technologies (FRQ-NT).

This work was supported by operating grants from the Canadian Institutes of Health Research (CIHR) PJ-155944 to JMDN and FDN–148372 to TM. PGS was partially supported by a fellowship from Fondation de recherche en Santé de Québec (FRQ-S) and the Cole Foundation. TM was supported by a Canada Research Chair (Tier 1) and JMDN by a Bourse de mérite from FRQ-S.

## SUPPLEMENTARY FIGURE LEGENDS

**Supplementary figure 1 – Characterization of mice with *Hnrnpl* deletion in B cells (Related to Figure 1)**

**A)** Body and spleen weights, **B)** splenic B and T cell counts, **C)** GC B cell counts, and **D)** representative flow cytometry plot and proportions of splenic NF, MZ, FO and GC B cell subpopulations in n *CD21-cre* (+/+), *CD21-cre Hnrnpl^F/+^*(F/+) and *CD21-cre Hnrnpl^F/F^* (F/F) mice. **E)** Body and spleen weights, and splenocyte counts, of lethally irradiated recipient mice that received bone marrow cells from *μMT* mice and either *CD21-cre Hnrnpl^F/+^* (F/+) or *CD21-cre Hnrnpl^F/F^* (F/F) mice. P-values are indicated if differences in group means are statistically significant (p<0.05) by one-way ANOVA with post-hoc Tukey’s multiple comparison test (B, C) or unpaired two-tailed t-test with Welch’s correction (for E).

**Supplementary figure 2 – Additional controls of competitive BM chimeras (Related to Figure 2)**

**A)** Initial proportion of CD45.1:CD45.2 BM cells transplanted into recipient mice. **B)** Cell counts and proportions of splenic B cells in recipient mice reconstituted with BM from CD45.1+ WT and either CD45.2+ *Rosa^mT/mG^ CD21-cre Hnrnpl^F/+^* (F/+) or *Rosa^mT/mG^ CD21-cre Hnrnpl^F/F^* (F/F). **C)** Representative flow cytometry plots and proportions of splenic newly formed B (NF), marginal zone B (MZ), follicular B (Fo) and germinal center (GC) B cells in F/+ and F/F mice. Results compiled from 2 independent experiments. Bars indicate mean ± SE of 8 mice per group. P-values are indicated for statistically significant differences (p<0.05) by unpaired, two-tailed Mann-Whitney test.

**Supplementary figure 3 – Cellular and transcriptional effects of hnRNPL loss in B cells (Related to Figure 3)**

Representative flow cytometry plots showing **A)** the proportions of hnRNPL-excised (GFP+; green) and non-excised (tdTomato+; orange) B cells and **B)** parameters indicating cell size (FSC) and granularity (SSC) of B cells, with quantitation for 4 mice, for splenic B cells from *Rosa^mT/mG^ CD21-cre Hnrnpl^F/+^*(F/+) or *Rosa^mT/mG^ CD21-cre Hnrnpl^F/F^* (F/F), before and after ex vivo activation with LPS and IL-4. **C, D)** Top 10 KEGG pathways enriched in upregulated (C) or downregulated (D) genes in hnRNPL-deficient versus WT cells. The dendrograms indicate degree of similarity (shared genes) between pathways. **E**) Quantification of gene expression by RT-qPCR for p21 (*Cdkn1a*), pro- and anti-apoptotic factors in resting (d0) or activated (48 h LPS + IL-4) (d2) B cells with from *CD21-cre Hnrnpl^F/+^*(F/+) or *CD21-cre Hnrnpl^F/F^* (F/F) mice. Results are compiled from n mice. Bars indicate mean ± SD. P-value summaries are indicated if differences in group means are statistically significant (p<0.05) by unpaired, two-tailed t-test with Welch’s correction (*p<0.05; ** p<0.01; *** p<0.001).

**Supplementary Figure 4 – Conserved roles of hnRNPL – (Related to Figure 5).**

**A)** Quantile-normalized expression levels of *HNRNPL* and *HNRNPLL* in the indicated control (CTL) or hnRNPL-depleted cells (left). Ratio of *HNRNPL* over *HNRNPLL* read counts in each control cell type. **B)** Quantile-normalized expression levels of indicated genes in the various control or hnRNPL-depleted cells. **C)** hnRNPL occupancy at the indicated loci in WT splenic B cells resting (d0) or activated with LPS + IL-4 for 2 days (d2 actB). **D)** Comparison of significantly up- and downregulated genes upon hnRNPL depletion shared among the indicated cell types. The insets show the relative expression of selected genes in the same cell types.

**Supplementary Figure 5 – Comparison of hnRNPL depletion effects among cell types (Related to Figure 6).**

**A)** Exon inclusion levels of an intermediate exon in selected genes from those listed in Fig 5B, in the indicated control (CTL) and hnRNPL-depleted cell types. The schemes indicate the positions of amino acids and/or stop codons coded by the included exon in the respective human genes. **B)** Sashimi plot showing the inclusion of an intermediate exon in *SIRT1* regulated by hnRNPL status in selected cell types. **C)** Quantile-normalized expression levels of indicated genes in the various CTL or hnRNPL-depleted cell types.

**Supplementary Figure 6 – Mitochondrial function-related gene expression (Related to Figure 7).**

**A)** Selected gene expression in splenic B cells activated ex vivo for 1 day with LPS + IL-4 from hnRNPL-deficient (GFP+ cells from *Rosa^mT/mG^ CD21-cre Hnrnpl^F/F^*) and WT (GFP+ cells from *Rosa^mT/mG^ CD21-cre*) mice. Individual independent samples (symbols) and means (bars) are plotted. **B)** Heatmap of selected genes from RNA seq described in Figure 4. **C)** Relative expression level of selected genes measured by RT-qPCR in resting (d0) and ex vivo LPS+IL-4-activated splenic B cells from *CD21-cre* (+/+) and *CD21-cre Hnrnpl^F/F^* (F/F) mice.

