## Supplementary figures for "A conserved role of hnRNPL in regulating alternative splicing of transcriptional regulators necessary for B cell activation"

1. Institut de Recherches Cliniques de Montréal, 110 avenue des Pins Ouest, Montréal, QC, H2W 1R7, Canada.
2. Department of Medicine, Division of Experimental Medicine, McGill University, 1001 Boulevard Decarie, Montreal, QC, H4A 3J1, Canada.
3. Molecular Biology Programs, Université de Montréal, C.P. 6128, succ. Centre-ville, Montréal, QC, H3C 3J7, Canada.
4. Département de microbiologie, infectiologie et immunologie, Université de Montréal, 2900 Boul Edouard-Montpetit, Montréal, QC, H3T 1J4, Canada.
5. Department of Medicine, Université de Montréal, C.P. 6128, succ. Centre-ville, Montréal, QC, H3C 3J7, Canada.
†Current address: Thermo Fisher Scientific Norway Holdings Ullernchausséen 52, 0379 Oslo, Norway

### SUPPLEMENTARY MATERIAL

- Supplementary Figure S1 – Characterization of mice with Hnrnpl deletion in B cells (related to Figure 1)
- Supplementary Figure S2 – Additional controls of competitive BM chimeras (related to Figure 2)
- Supplementary Figure S3 – Cellular and transcriptional effects of hnRNPL loss in B cells (related to Figure 3)
- Supplementary Figure S4 – Conserved roles for hnRNPL in mouse and human cells (related to Figure 5)
- Supplementary Figure S5 – Splicing effects of hnRNPL loss among cell types (related to Figure 6)
- Supplementary Figure S6 – Mitochondrial function-related gene expression (related to Figure 7)
- File with uncropped western blots and agarose gels.

Supplementary resources (Provided as separate excel files)

- Supplementary Table S1 – Differentially expressed genes in *Hnrnpl-/-* vs control activated B cells
- Supplementary Table S2 – Comparative analysis of differentially expressed genes in various hnRNPL-deficient vs WT cell types
- Supplementary Table S3 – Functional annotation of genes bearing splicing changes in hnRNPL-deficient B cells
- Supplementary Table S4 – Comparative analysis of splicing changes across various hnRNPL-deficient vs WT cell types
- Supplementary Table S5 – Antibodies
- Supplementary Table S6 – Oligonucleotides

Supplementary figure 1 – Characterization of mice with Hnrnpl deletion in B cells (Related to Figure 1)

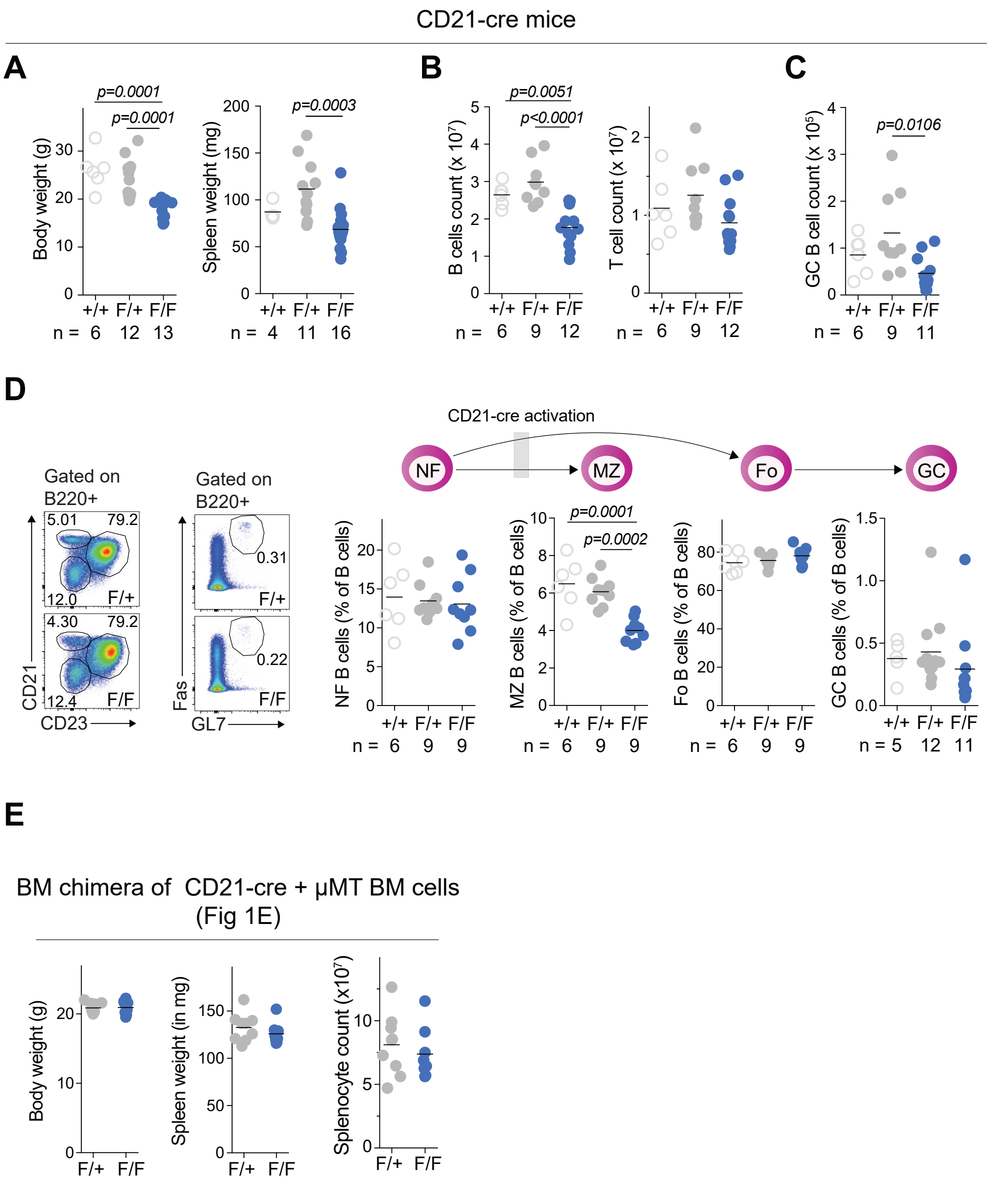


A) Body and spleen weights,
B) splenic B and T cell counts,
C) GC B cell counts, and
D) representative flow cytometry plot and proportions of splenic NF, MZ, FO and GC B cell subpopulations in n CD21-cre (+/+), CD21-cre HnrnplF/+ (F/+) and CD21-cre HnrnplF/F (F/F) mice.
E) Body and spleen weights, and splenocyte counts, of lethally irradiated recipient mice that received bone marrow cells from µMT mice and either CD21-cre HnrnplF/+ (F/+) or CD21-cre HnrnplF/F (F/F) mice.
P-values are indicated if differences in group means are statistically significant ($p<0.05$) by one-way ANOVA with post-hoc Tukey's multiple comparison test (B, C) or unpaired two-tailed t-test with Welch's correction (for E).

Supplementary figure 2 – Additional controls of competitive BM chimeras (Related to Figure 2)

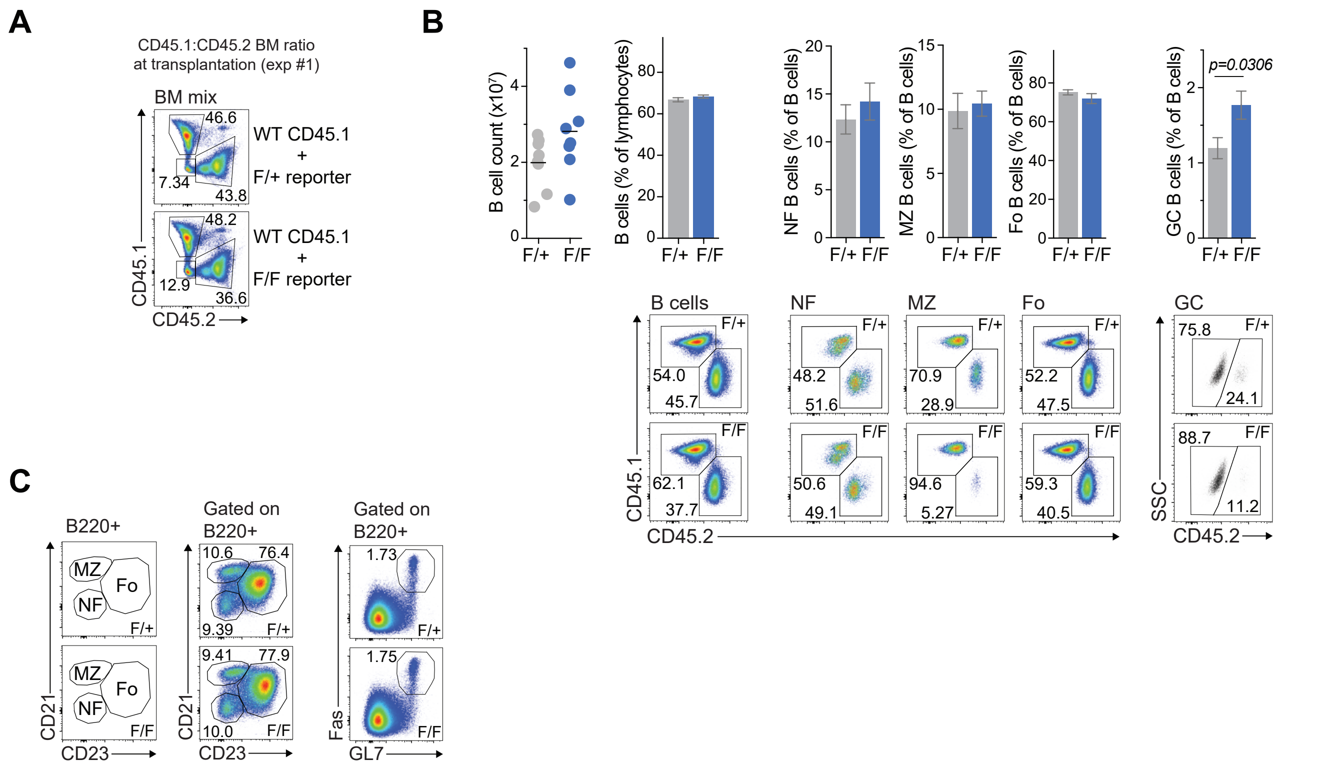


A) Initial proportion of CD45.1:CD45.2 BM cells transplanted into recipient mice.
B) Cell counts and proportions of splenic B cells in recipient mice reconstituted with BM from CD45.1+ WT and either CD45.2+ RosamT/mG CD21-cre HnrnplF/+ (F/+) or RosamT/mG CD21-cre HnrnplF/F (F/F).
C) Representative flow cytometry plots and proportions of splenic newly formed B (NF), marginal zone B (MZ), follicular B (Fo) and germinal center (GC) B cells in F/+ and F/F mice. Results compiled from 2 independent experiments. Bars indicate mean ± SE of 8 mice per group. P-values are indicated for statistically significant differences ($p<0.05$) by unpaired, two-tailed Mann-Whitney test.

Supplementary figure 3 – Cellular and transcriptional effects of hnRNPL loss in B cells (Related to Figure 3)

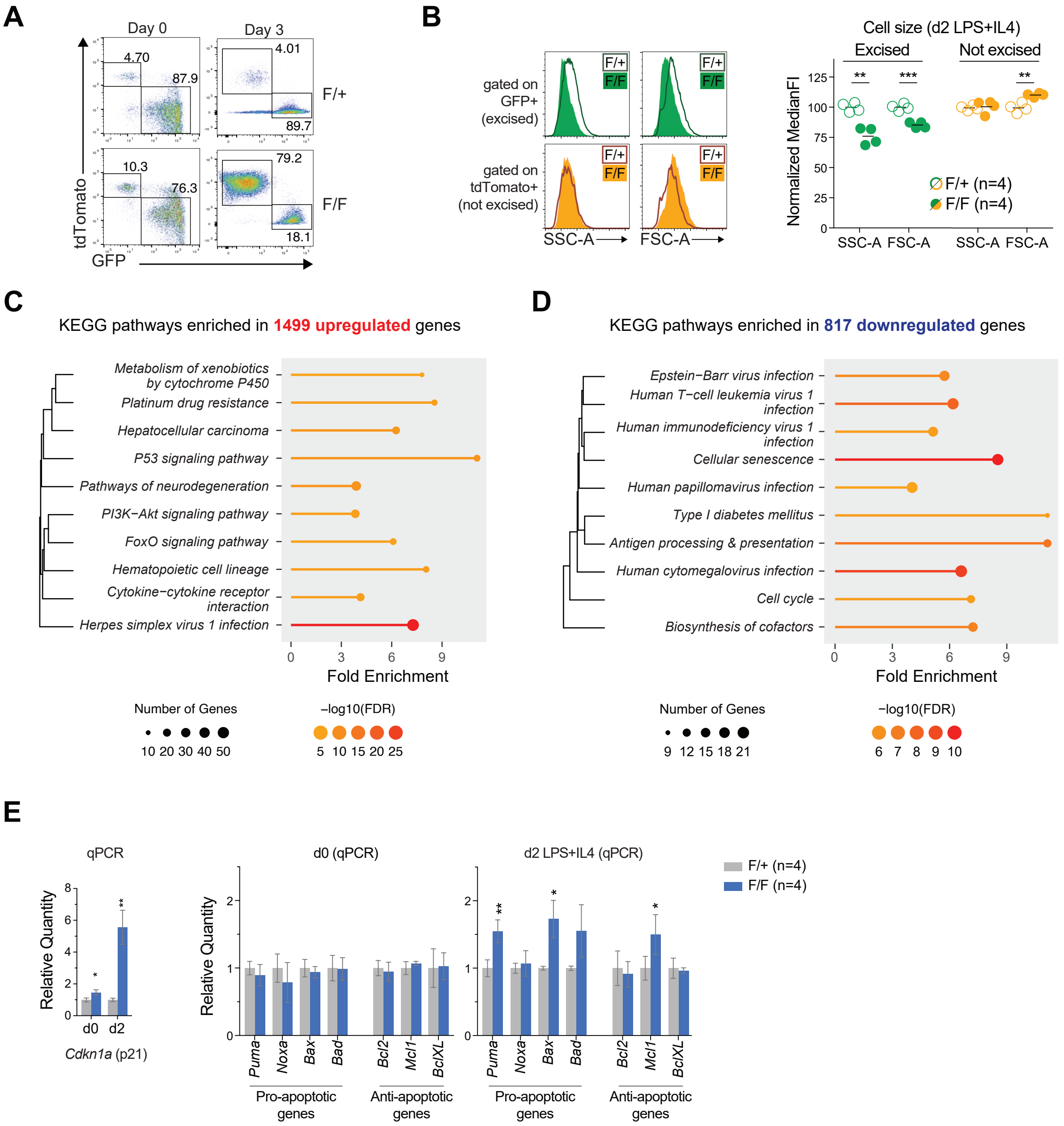


Representative flow cytometry plots showing A) the proportions of hnRNPL-excised (GFP+; green) and non-excised (tdTomato+; orange) B cells and
B) parameters indicating cell size (FSC) and granularity (SSC) of B cells, with quantitation for n mice, for splenic B cells from RosamT/mG CD21-cre hnRNPLF/+ (F/+) or RosamT/mG CD21-cre HnrnplF/F (F/F), before and after ex vivo activation with LPS and IL-4.
C, D) Top 10 KEGG pathways enriched in upregulated or downregulated genes in hnRNPL-deficient versus WT cells. The dendrograms indicate degree of similarity (shared genes) between pathways.
E) Quantification of gene expression by RT-qPCR for p21 (Cdkn1a), pro- and anti-apoptotic factors in resting (d0) or activated (48 h LPS + IL-4) (d2) B cells with from CD21-cre (+/+) or CD21-cre HnrnplF/F (F/F) mice. Results are compiled from n mice. Bars indicate mean ± SD. P-value summaries are indicated if differences in group means are statistically significant ($p<0.05$) by unpaired, two-tailed t-test with Welch’s correction (*$p<0.05$; ** $p<0.01$; *** $p<0.001$).

Figure S4: Conserved roles for hnRNPL in mouse and human cells (Related to Figure 5)

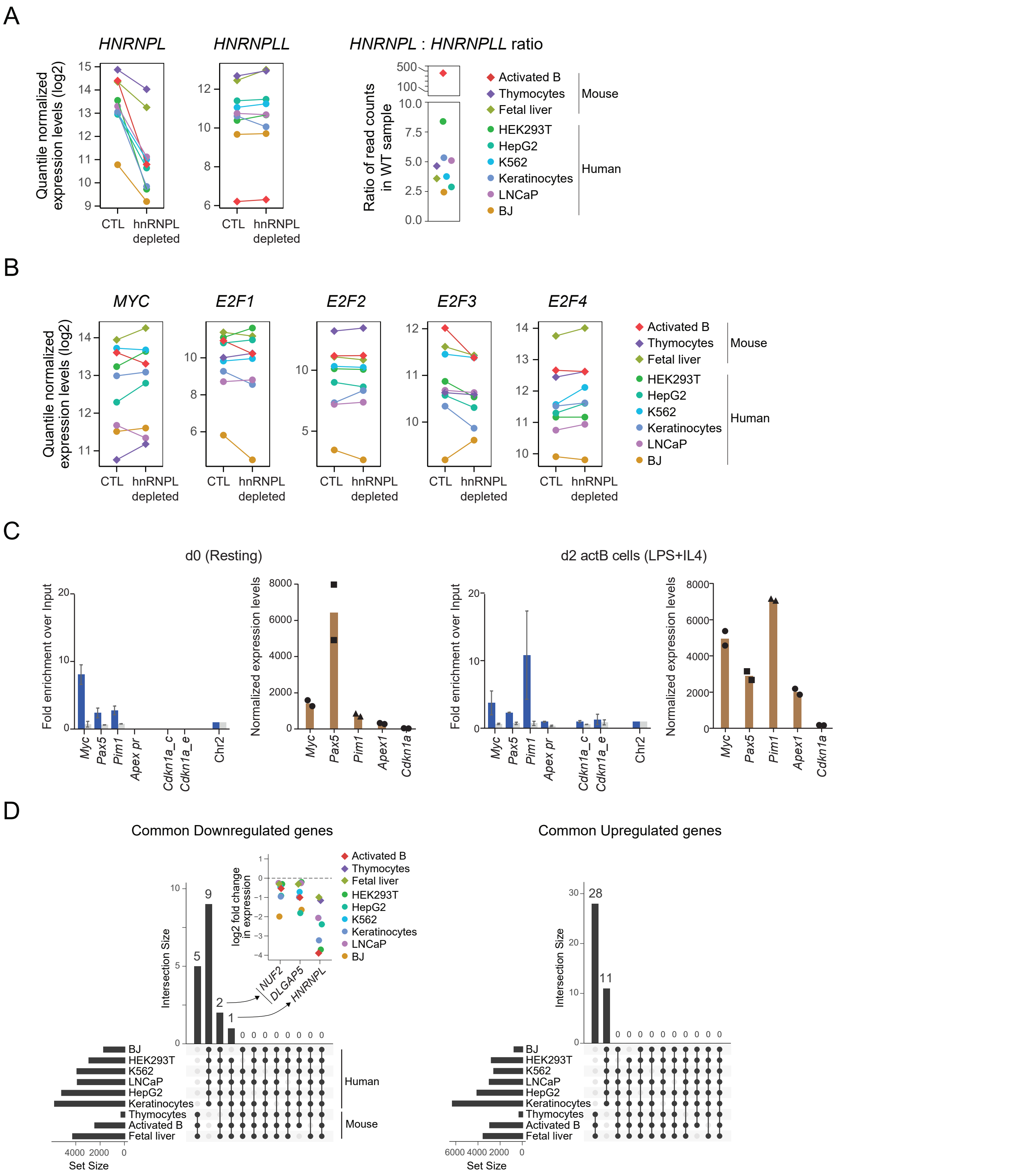


A) Quantile-normalized expression levels of HNRNPL and HNRNPLL in the indicated control (CTL) or hnRNPL-depleted cells (left). Ratio of HNRNPL over HNRNPLL read counts in each control cell type. B) Quantile-normalized expression levels of indicated genes in the various control or hnRNPL-depleted cells. C) hnRNPL occupancy at the indicated loci in WT splenic B cells resting (d0) or activated with LPS + IL-4 for 2 days (d2 actB). D) Comparison of significantly up- and downregulated genes upon hnRNPL depletion shared among the indicated cell types. The insets show the relative expression of selected genes in the same cell types.

Supplementary Figure 5 – Splicing effects of hnRNPL loss among cell types (Related to Figure 6).

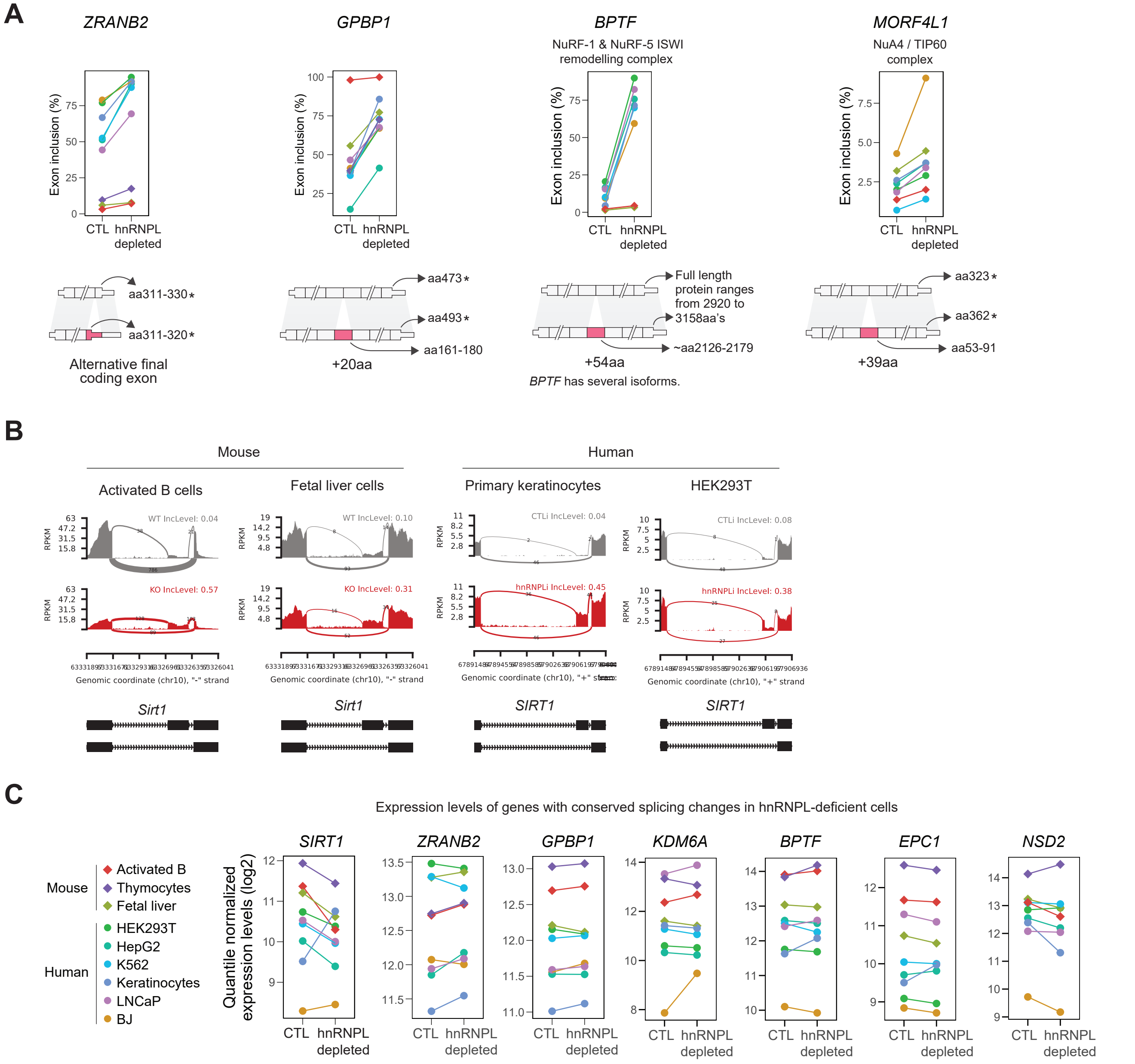


A) Exon inclusion levels of an intermediate exon in selected genes from those listed in Fig 5B, in the indicated control (CTL) and hnRNPL-depleted cell types. The schemes indicate the positions of amino acids and/or stop codons coded by the included exon in the respective human genes.
B) Sashimi plot showing the inclusion of an intermediate exon in SIRT1 regulated by hnRNPL status in selected cell types.
C) Quantile-normalized expression levels of indicated genes in the various CTL or hnRNPL-depleted cell types.

Supplementary Figure 6 – Mitochondrial function-related gene expression (related to Figure 7)

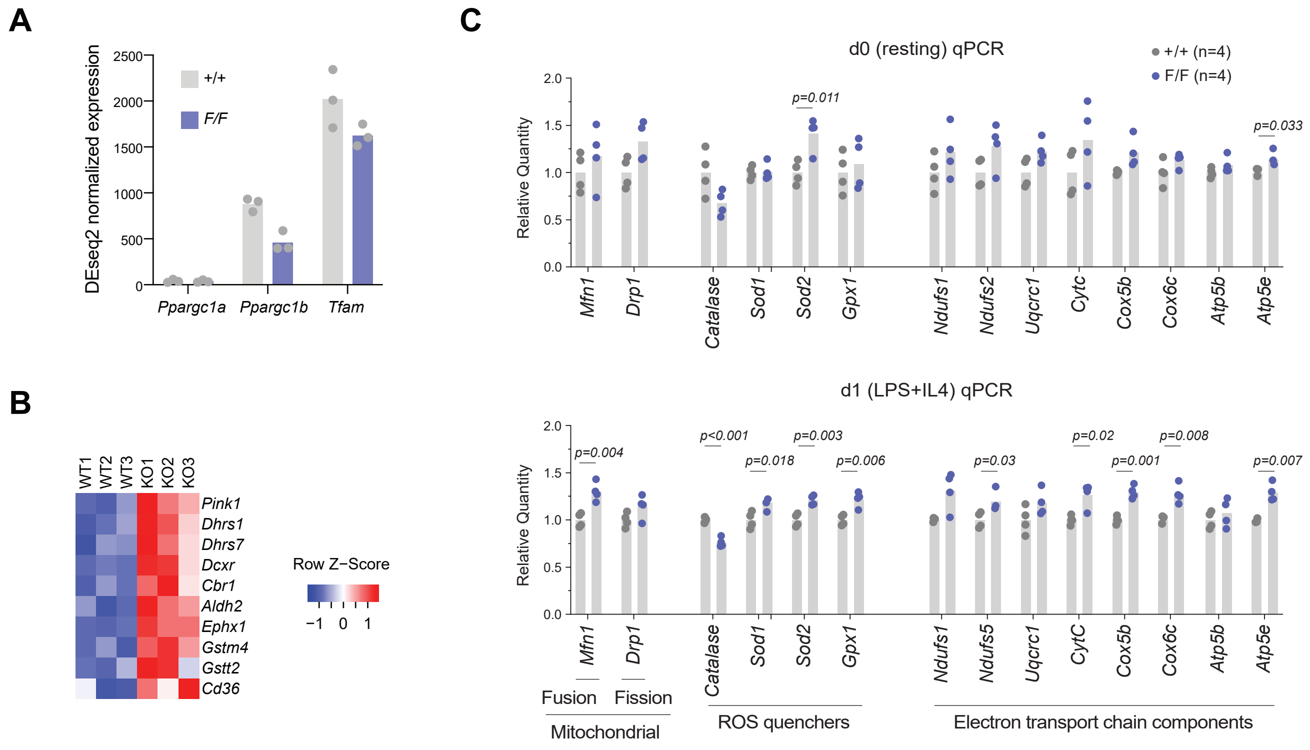


A) Selected gene expression in splenic B cells activated ex vivo for 1 day with LPS + IL-4 from hnRNPL-deficient (GFP+ cells from RosamT/mG CD21-cre HnrnplF/F) and WT (GFP+ cells from RosamT/mG CD21-cre) mice. Individual independent samples (symbols) and means (bars) are plotted.
B) Heatmap of selected genes from RNA seq described in Figure 4.
C) Relative expression level of selected genes measured by RT-qPCR in resting (d0) and ex vivo LPS+IL-4-activated splenic B cells from CD21-cre (+/+) and CD21-cre HnrnplF/F (F/F) mice.

Figure 1J

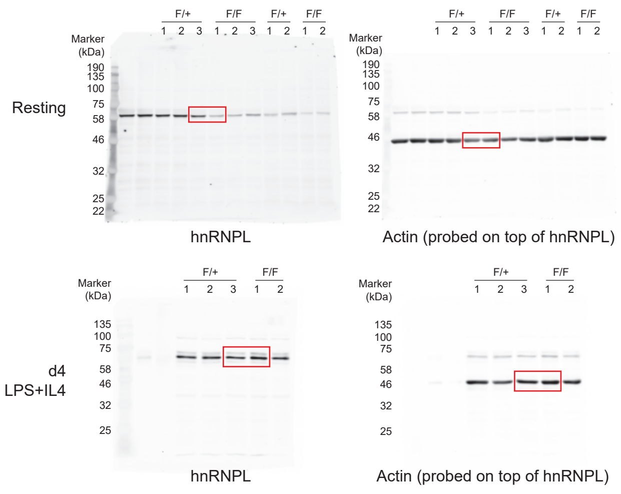


Figure 5F

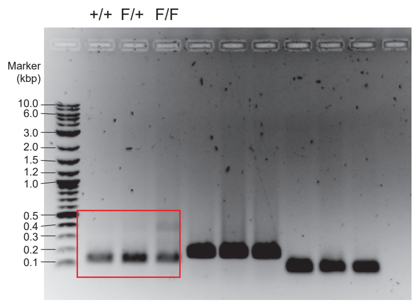
